# An *in situ* atlas of mitochondrial DNA in mammalian tissues reveals high content in stem/progenitor cells

**DOI:** 10.1101/2019.12.19.876144

**Authors:** Jiayu Chen, Qizhi Zheng, Lauren B. Peiffer, Jessica L. Hicks, Michael C. Haffner, Avi Z. Rosenberg, Moshe Levi, Xiaoxin X. Wang, Busra Ozbek, Srinivasan Yegnasubramanian, Angelo M. De Marzo

## Abstract

Mitochondria regulate ATP production, metabolism and cell death. Alterations in mitochondrial DNA (mtDNA) sequence and copy number are implicated in aging and organ dysfunction in diverse inherited and sporadic diseases. Since most measurements of mtDNA use homogenates of complex tissues, little is known about cell type-specific mtDNA copy number heterogeneity in normal physiology, aging and disease. Thus, the precise cell types whose loss of mitochondrial activity and altered mtDNA copy number that result in organ dysfunction in aging and disease have often not been clarified. Here, we validated an *in situ* hybridization approach to generate a single cell resolution atlas of mtDNA content in mammalian tissues. In hierarchically organized self-renewing tissues, higher levels of mtDNA were observed in stem/proliferative compartments compared to differentiated compartments. Striking zonal patterns of mtDNA levels in the liver reflected the known oxygen tension gradient. In the kidney, proximal and distal tubules had markedly higher mtDNA levels compared to cells within glomeruli and collecting duct epithelial cells. Decreased mtDNA levels were visualized in renal tubules as a function of aging, which was prevented by calorie restriction. We provide a novel approach for quantifying species- and cell type-specific mtDNA copy number and dynamics in any normal or diseased tissue and can be used for monitoring the effects of interventions in animal and human studies.

## INTRODUCTION

Mitochondria are critical for many cellular functions including ATP production, metabolism, macromolecular synthesis, calcium homeostasis, cell signaling and programmed cell death.^1^ Mitochondria have their own genome (mitochondrial DNA or mtDNA), encoding 16 proteins involved in oxidative phosphorylation, 22 transfer RNAs, and 2 ribosomal RNAs required for mitochondrial ribosome function. Over the last few decades a large number of genetic diseases of mitochondrial function have been characterized that are caused by maternally inherited mutations in the mitochondrial genome, or by nuclear encoded genes whose products function in mitochondria.^1, 2^ These mutations often result in disorders of the nervous system (e.g. epilepsy, deafness, neuropathy, ataxia), skeletal muscle (e.g. weakness, ophthalmoplegia), and cardiac muscle (e.g. cardiomyopathy). Interestingly, a number of inherited diseases caused by mtDNA mutations, or mutations in nuclear genes whose products function in mitochondria, demonstrate organ-specific and cell type-specific phenotypes.^3^

Mitochondrial dysfunction can also be acquired in the absence of inherited mutations, and has been linked to declines of function with aging in several organ systems and to age-related cardiovascular and neurodegenerative diseases, as well as cancer.^4–7^ Whether these age-related declines in mitochondrial function directly result from the known accumulation of somatic mtDNA mutations, or sub-genomic deletions with age, is still unclear.^6, 8^

Mitochondrial function can be regulated through mitochondrial biogenesis, which is controlled in a cell-type and tissue-specific manner and is determined in part by regulating mtDNA copy number.^9^ As such, there has been interest in measuring mtDNA copy number (independent of mtDNA mutations) in aging and diseased tissues.^10–16^ In human studies, while there is general agreement that mtDNA copy number declines with age in peripheral blood mononuclear cells,^17–20^ no consensus has been reached regarding mtDNA copy number changes with aging in non-blood related tissue types.^15^ This is the case even in skeletal muscle, in which both overall cellular/organ function and mitochondrial function are well-known to decline with age. The discrepancies in the results from these studies do not appear to be explained simply by methodological differences.^15^ Since widely employed strategies for measuring mtDNA levels are performed using solution-based methods after cellular disruption/homogenization of bulk tissues, the measurements necessarily reflect a composite of heterogeneous cell types. Therefore, it has been suggested that some of the discrepancies may be related to the fact that tissues are composed of multiple cell types, whose proportions can change in a variable manner with aging due to processes such as inflammation, regeneration, and fibrosis.^8^ One method that has been used in attempts to circumvent the problem of heterogeneous cell types is to isolate specific cell populations (e.g. in the liver and CNS) by laser capture microdissection followed by quantitative PCR (qPCR) or genomic sequencing.^16, 21^ Yet, even using laser capture it can be very difficult to evaluate specific cell types or single cells across tissue samples. Therefore, an *in situ* method for mtDNA measurements would be advantageous. While prior studies have used *in situ* approaches to localize mtDNA in cells and tissues, these methods were used in limited applications, such as those regarding mtDNA replication and transcription, as well as to examine mtDNA levels in specific cell populations in a single tissue type or disease state.^22–33^ In this study we developed a novel quantitative *in situ* assay to interrogate mtDNA at the single cell level in formalin fixed paraffin embedded (FFPE) or frozen archival tissue. By contrast to bulk methods, this *in situ* approach can be implemented using standard bright field microscopy facilitating rapid spatial determination of mtDNA content at single cell resolution in complex tissues.

## MATERIALS AND METHODS

### Cell Culture and Treatment

Prostate cancer cell lines CWR22Rv1 (CRL-2505), LNCaP (CRL-1740) and PC3 (CRL-1435) were obtained from American Type Culture Collection (ATCC, Manassas, VA). All cell lines were cultured in RPMI 1640 with L-glutamine media (Corning, Corning, NY) and 10% heat inactivated BenchMark™ Fetal Bovine Serum (FBS, Gemini Bio-Products, West Sacramento, CA). Cells were maintained in 75 cm^2^ or 175 cm^2^ TC flasks (Sarstedt, Germany) in a humidified incubator at 37°C with 5% CO_2_. When reaching ∼85% confluency, cells were passaged using TrypLE™ Express Enzyme without phenol red (Thermo Fisher, Waltham, MA). All cell lines were authenticated using Short Tandem Repeat (STR) profiling by the Genetic Resources Core Facility at The Johns Hopkins University School of Medicine.

To generate cells with different level of mtDNA, CWR22Rv1 and PC3 cells were cultured in RPMI 1640 with L-glutamine media (Corning) containing 2’,3’-Dideoxycytidine (ddC, D5782, MilliporeSigma, Burlington, MA), 10% FBS, 1 mM Sodium Pyruvate (Thermofisher) and 50 μg/mL Uridine (MilliporeSigma). ddC containing media was replaced twice a week.

### Cell Block Preparation

Formalin-fixed paraffin-embedded (FFPE) cell blocks were generated using the protocols described previously with the following modifications.^34^ Cells were harvested by trypsinization as above, washed once with PBS, and resuspended in ∼ 150 μL 10% neutral buffered formalin (NBF). The resuspended cells were transferred to a 0.5 mL microcentrifuge tube with solidified 2% agarose prefilled in the tapered portion of the bottom, and spun down in a swinging bucket centrifuge at 1000 rpm for 5 min. The resulting supernatant was removed, and fresh 10% NBF was added without disturbing the pellet. The microcentrifuge tube was spun at 1500 rpm for 5 min, and the resulting cell plug was fixed in formalin for 48 hours at room temperature by submerging the microcentrifuge tube into a 15 mL conical tube containing 10% NBF. After fixation, the microcentrifuge tubes were transferred to a new 15 mL conical tube with PBS and stored at 4°C prior to processing and embedding in paraffin blocks.

### Human Prostate, Small Intestine, Liver, Kidney and Tissues

Prostate tissues were obtained from radical prostatectomies for prostate cancer. Human duodenum (small intestine) was obtained as frozen tissue samples of grossly normal duodenum from patients undergoing pancreatoduodenectomy for pancreatic cancer. Human liver and kidney tissues were obtained from samples of patients undergoing autopsy for metastatic prostate cancer. This study was approved by the Johns Hopkins institutional internal review board.

### Normal Mouse Tissues

Experimentally unmanipulated wild type normal (FVB/N mice (derived from FVB-Tg (ARR2/Pbsn-MYC)7Key/Nci) were conventionally housed according to protocols approved by the Animal Care and Use Committee at Johns Hopkins University. Animals (2-3 months, n = 3 for each sex) were euthanized, necropsied dissected as previously described,^35^ and tissues were fixed in 10% NBF or Formical-2000 for bone tissues samples (StatLab) for 48 hours and processed into FFPE blocks as above.

### Mouse Aging and Calorie Restriction Study

Male mice (C57BL/6, NIA) were housed and maintained on a 12 hour light-dark cycle, at a constant temperature (22 ± 2°C) and relative humidity (55 ± 15%) and treated as described.^36^ Mice were randomly divided into the following groups: Group 1 - control young (6 months, n=3); Group 2 - control aging (24 months, n=3), Group 3 - aging with calorie restriction (24 months, n=3). Tap water was available ad libitum and rodent diets were available *ad libitum* for groups 1 and 2. For group 3, mice were calorie restricted (40% calorie reduction) from 14 weeks of age. Immediately following euthanasia, the kidneys were harvested and processed for FFPE.

### mtDNA ISH Assay

Chromogenic *in situ* hybridization (CISH) of mtDNA on FFPE prostate tissues and FFPE cell blocks was performed manually following the instructions from the manufacturer (Advanced Cell Diagnostics, or ACD, Newark, CA) using RNAscope® 2.5 HD Detection Kit (BROWN) with the following modifications.^37^ Slides were baked at 60°C for 30 min, deparaffinized in xylene 3 times for a total of 20 min, and in 100% ethanol (EtOH) 2 times for 4 min. Slides were then incubated in Pretreatment I solution (H_2_O_2_) for 10 min at room temperature, and steamed in Pretreatment II solution for 18 min. Slides were dipped in deionized water (diH_2_O) with 0.1% Tween 20 (Source) once, and incubated in Protease Plus solution at 40°C as Pretreatment III. The protease digestion time depended on the sample types and species from 10 min (human cell blocks) to 20 or 30 min for the human and mouse tissues. Slides were incubated in Hs-MT-COX1-sense probe (Cat No. 478051) or Mm-Mt-Cox1-sense probe (Cat No. 530891), depending on the species, for 2 hours followed by the standard amplification steps as instructed by the manufacturer. Probes were diluted in RNAscope® Probe Diluent and used at 1:200 for FFPE human prostate tissues and cell blocks, and 1:300 for FFPE mouse tissues.

Whole slides were scanned using a Roche-Ventana DP200 (Roche) or a Hamamatsu Nanozoomer XR whole slide scanner (Hamamatsu Photonics, Japan), and uploaded to Concentriq® (Proscia, Philadelphia PA) for whole slide image viewing and micrograph documentation.

For HALO image analysis, the Hs-MT-COX1-sense probe was used at 1:200 dilution, and visualized by alcohol-soluble ImmPACT® AEC Peroxidase (HRP) Substrate (SK-4205, Vector Laboratory, Burlingame, CA), mounted in aqueous media (VectaMount® AQ, H-5501, Vector Laboratory). After whole slide scanning, coverslips were removed by soaking in diH_2_O and residual mounting media was washed away with fresh diH_2_O. The polymerized AEC chromogenic situ signals were then removed by washing in 100% ethanol, and these “hematoxylin-only” slides were mounted in Cytoseal 60 (8310-16, Thermo Fisher) and scanned again. Under these conditions, hematoxylin stains the entire cell including the cytoplasm. The ratio of the area of AEC-positive signals to total cellular area was used as the area-fraction of mtDNA signals.

### ISH Validation

DNase and RNase pretreatments were performed using the manufacturer’s suggested protocol (ACD) with the following modifications. For DNase pretreatment, after steaming in Pretreatment II, the slides were treated with 100 μL DNase reaction buffer containing 10 μL DNase I Reaction Buffer (10X), 1 μL (2 units) DNAse I (M0303S, New England BioLabs, Ipswich, MA), and 89 μL nuclease-free H_2_O. The slides were then incubated at 37°C for 10 min, washed in diH_2_O for 3 times, followed by *in situ* Pretreatment III. Similarly, for RNase pretreatment, the slides were immersed in 50 μg/mL RNase A (EN0531, ThermoFisher) diluted in 0.01 M Tris-HCl buffer, incubated at 37°C for 30 min, washed extensively in phosphate buffered saline with Tween-20 (PBST, P3563, MilliporeSigma) followed by *in situ* Pretreatment II. Human MT-COX1-sense probe was used at 1:50 dilution. Xenografts of human PC3 and VCaP prostate cancer cell lines were performed as described previously.^38^ Detection of mtDNA on mouse xenograft slides was performed using the CISH-DAB protocol as mentioned above. Both Mouse MT-COX1-sense probe and human MT-COX1-sense probe were used at 1:50 dilution.

Fluorescence *in situ* hybridization (FISH) of mtDNA on FFPE tissues was conducted manually according to the manufacturer’s protocols using RNAscope® Multiplex Fluorescent Kit v2 (ACD).^37^ Pretreatment and probe hybridization procedures remained the same as the one mentioned above for standard mtDNA CISH. During the signal amplification and development process, Opal™ Dyes (NEL741E001KT, NEL744E001KT and NEL744E001KT, PerkinElmer, Waltham, MA) were used at 1:3500 in the TSA buffer provided in the RNAscope® Multiplex Fluorescent Kit v2 for each probe. The slides were counterstained with DAPI, mounted in ProLong™ Gold Antifade Mountant (Thermo Fisher) and coverslipped.

Cultured adherent cells were prepared for RNAscope® *in situ* hybridization using the manufacturer’s instructions with following modifications. CWR22Rv1 and PC3 cells were seeded on Nunc™ Lab-Tek™ II CC2™ Chamber Slides (Thermo Fisher) at ∼ 30,000 cells/cm^2^ two days before fixation. Cells were fixed for 30 min in 10% NBF, treated with H_2_O_2_, steamed in Pretreatment II solution for 5 min, and incubated in Protease III diluted 1:15 in PBS before proceeding to probe hybridization.

Localization of the mtDNA *in situ* signal was validated using a multiplex assay consisting of FISH for mtDNA and immunofluorescence for a mitochondrial protein MRPL12.^39, 40^ Briefly, LNCaP cells grown on chamber slides were fixed in formalin and mtDNA FISH was performed using the same protocol as for FFPE tissues. The slides were incubated in MRPL12 primary antibody (HPA022853, MilliporeSigma) at 1:200 dilution in Antibody Dilution Buffer (ADB250, Roche, Switzerland) for 45 min at RT. Alexa Fluor 568 Goat anti-Rabbit IgG (H+L) antibody (A-11036, Thermo Fisher) was used as secondary antibody at 1:300 dilution in PBS for 45 minutes at room temperature. The slides were counterstained, mounted, and coverslipped in the same fashion as mentioned above for FISH. Confocal fluorescent images were taken using a Zeiss AxioObserver inverted microscope with LSM700 confocal module (Carl Zeiss, Germany).

### Immunohistochemistry

Chromogenic immunohistochemistry was performed mainly using the automated Ventana Discovery ULTRA (Roche). The slides were steamed for 32 min in CC1 buffer (950-124, Roche) and incubated in the corresponding primary antibody at room temperature for 36 min. Anti-MTCO1 antibody was used at 1:8000 dilution (ab14705, abcam, Cambridge, MA) and Anti-COX IV antibody was used at 1:8000 dilution (3E11, 4850, Cell Signaling, Danvers, MA) as primary antibodies. The slides were developed using Discovery HQ HRP hapten-linked multimer detection kit (Roche).

### Quantitative PCR

Total DNA from cell lines was extracted using the DNeasy Blood & Tissue Kit (Qiagen, Germany). Quantitative PCR was performed using TaqMan™ Universal Master Mix II, no UNG (Catalog # 4440043) and TaqMan® Gene Expression Assays (Thermo Fisher) on a CFX Connect™ or a CFX96™ Real-Time PCR Detection System (BioRad, Hercules, CA). The cycling conditions were: 95°C for 10 min, 40 cycles of 95°C for 15 seconds, 60°C for 1 min, followed by melt curve analysis. The quantity of each amplified product was calculated using a standard curve of genomic DNA from peripheral blood leukocytes from a healthy donor (Catalog #: D1234148, Lot #: C210220, Biochain, Newark, CA). To calculate relative mtDNA copy number, MT-CO1 and MT-ND1 genes were used as specific mitochondrial targets, and the beta globin locus transcript 3 (BGLT3) and complement 2 (C2) genes were selected to be the endogenous controls. Taqman® Assays were purchased from Thermo Fishers. Information was as following: *MT-CO1*, Assay ID Hs02596864_g1, Catalog # 4331182; *MT-ND1*; *BGLT3*, Assay ID Hs01629437_s1, Catalog # 4331182; and *C2*, Assay ID Hs03599683_cn, Catalog # 4400291. All qPCR reactions were performed in triplicate.

## RESULTS

### Development and Validation of an in situ mtDNA Copy Number Assay

Transcription of the circular mtDNA proceeds in both directions. Therefore, when targeting mtDNA by *in situ* hybridization, it is possible that probes will hybridize to either RNA or DNA, resulting in ambiguity in signal interpretation. However, most light strand transcripts are rapidly degraded, including those anti-sense to the *MT-CO1* gene.^41, 42^ Thus, to minimize the potential of hybridization to mitochondrial RNA (mtRNA), we designed an ACD probe set targeting the antisense region of the *MT-CO1* gene (probe set referred to as Sense-MT-CO1-A). Figure 1A shows the results of chromogenic *in situ* hybridization (DNAscope CISH) using this probe set, in which highly intense DAB signals localize in a punctate distribution within the cytoplasm of an FFPE sample of human prostate cancer. Pre-treatment of slides with DNase I, but not with RNase A, abolished the signal, confirming specificity for DNA (Figure 1). To further validate the assay specificity, we treated prostate cancer cells (CWR22Rv1 and PC3) with 2′,3′-Dideoxycytidine (ddC) to reduce global mtDNA levels.^43^ The nucleotide derivative of ddC, ddCTP, can interact with and inhibit the activity of DNA polymerase gamma (pol γ), which is the only DNA polymerase in the mitochondria, and thus inhibit mtDNA replication.^43^ Figure 2 shows the results in which there was a ddC dose responsive reduction in hybridization signals in both cell lines.

**Figure 1.**
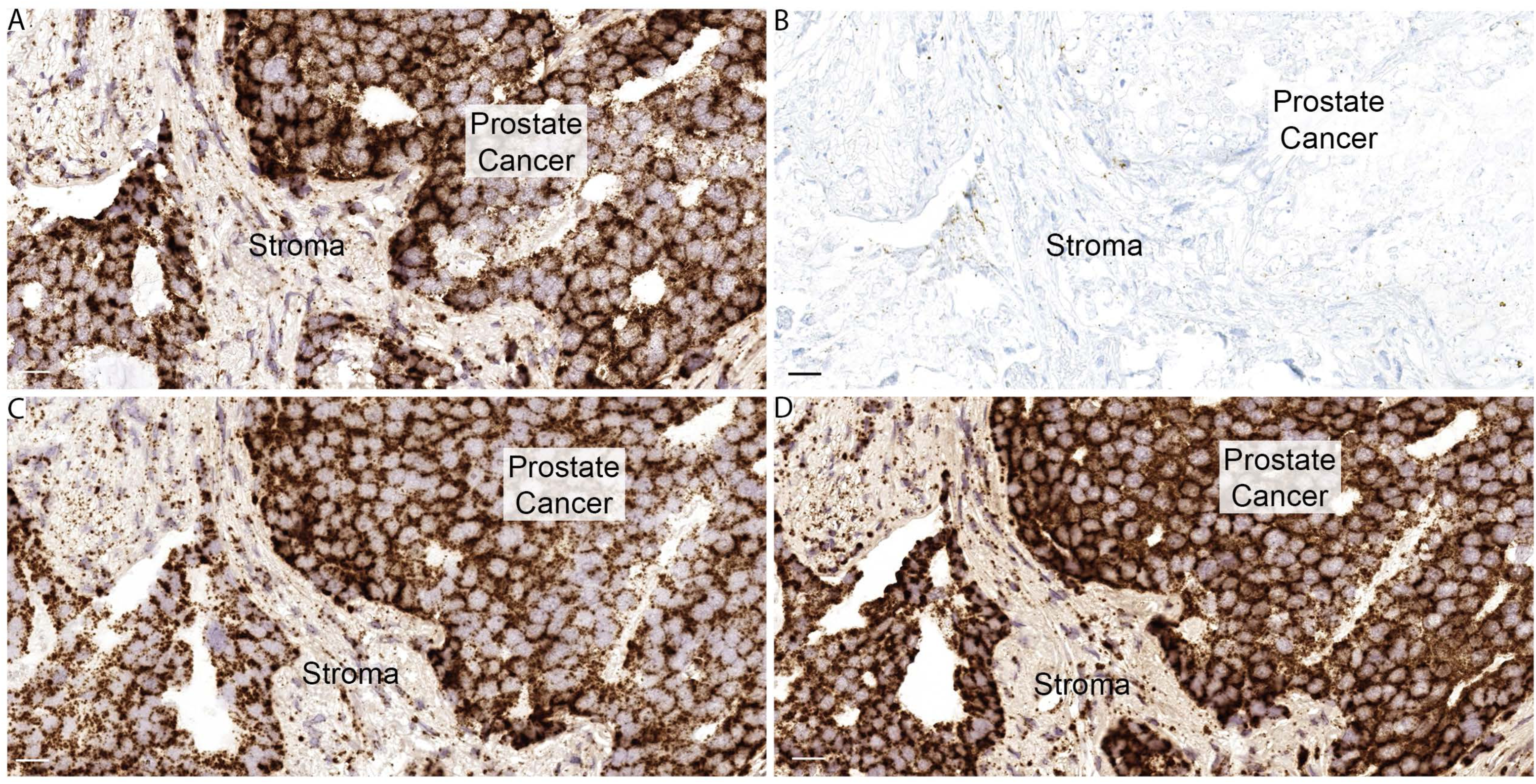
*In situ* assay specificity verified by DNase pretreatment. *MT-CO1* sense DNA *in situ* hybridization assay on FFPE prostate tissues without RNase A (**A**), with RNase A (**B**), without DNase I (**C**), and with DNase I (**D**) pretreatments. Original magnification x40.

**Figure 2.**
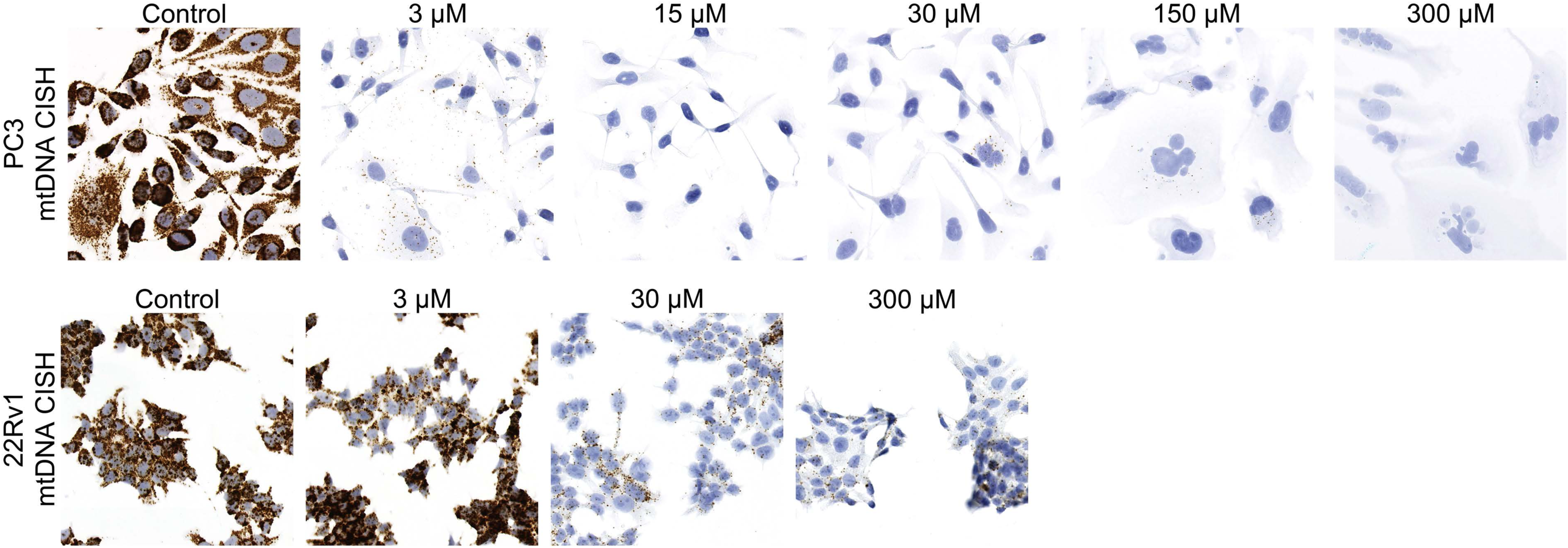
Dose-dependent stepwise reduction in hybridization signals after forced reductions in mtDNA by 2′,3′-Dideoxycytidine (ddC) in (**A**) PC3 and (**B**) 22Rv1 prostate cancer cell lines. The cells were treated for 9 days, and fixed in 10% formalin and processed into paraffin blocks and sectioned. Original magnification x20.

To determine if this *in situ* assay would provide a truly quantitative assessment of the mtDNA content, we correlated the mtDNA levels measured by CISH with mtDNA measurements by quantitative PCR (qPCR).^44, 45^ As standards, we treated PC3 and CWR22Rv1 cells with a single dose of ddC for varying times. At each time point, cells were harvested and split into two portions. The first was prepared for formalin fixation and paraffin embedding (FFPE), and the second was used for qPCR using the *MT-CO1* mtDNA locus normalized to the nuclear DNA gene, *BGLT3*. In a second qPCR reaction we used the mtDNA encoded *MT-ND1* gene that was normalized to a different nuclear gene (*C2)*. Figure 3A shows a stepwise reduction in the ratio of mtDNA to nuclear DNA for each primer/probe set used for qPCR after treating cells for increasing times with ddC. The ratios of mtDNA/nuclear DNA content from these two sets of primer pairs were highly correlated (Figure 3B). Figure 4A shows a corresponding stepwise decrease in mtDNA *in situ* signals in relation to duration of drug treatment. While both cell lines showed sharp reductions in mtDNA by both methods, in PC3 cells treated for 9 days the mtDNA signals were largely undetectable by CISH (Figure 4A) and extremely low by qPCR (Figure 3). For image analysis, we performed CISH using the alcohol soluble chromogenic substrate, AEC (Figure 4B). After hybridization and slide scanning, the chromagen (AEC) was dissolved in EtOH and the slides were re-scanned using the hematoxylin signals only (Figure 4B**, upper and lower right**). The ratio of mtDNA area to total cellular area was determined using Halo™. There was a strong positive correlation between real-time qPCR and the area-fraction of mtDNA signals by image analysis (Figure 4C).

**Figure 3.**
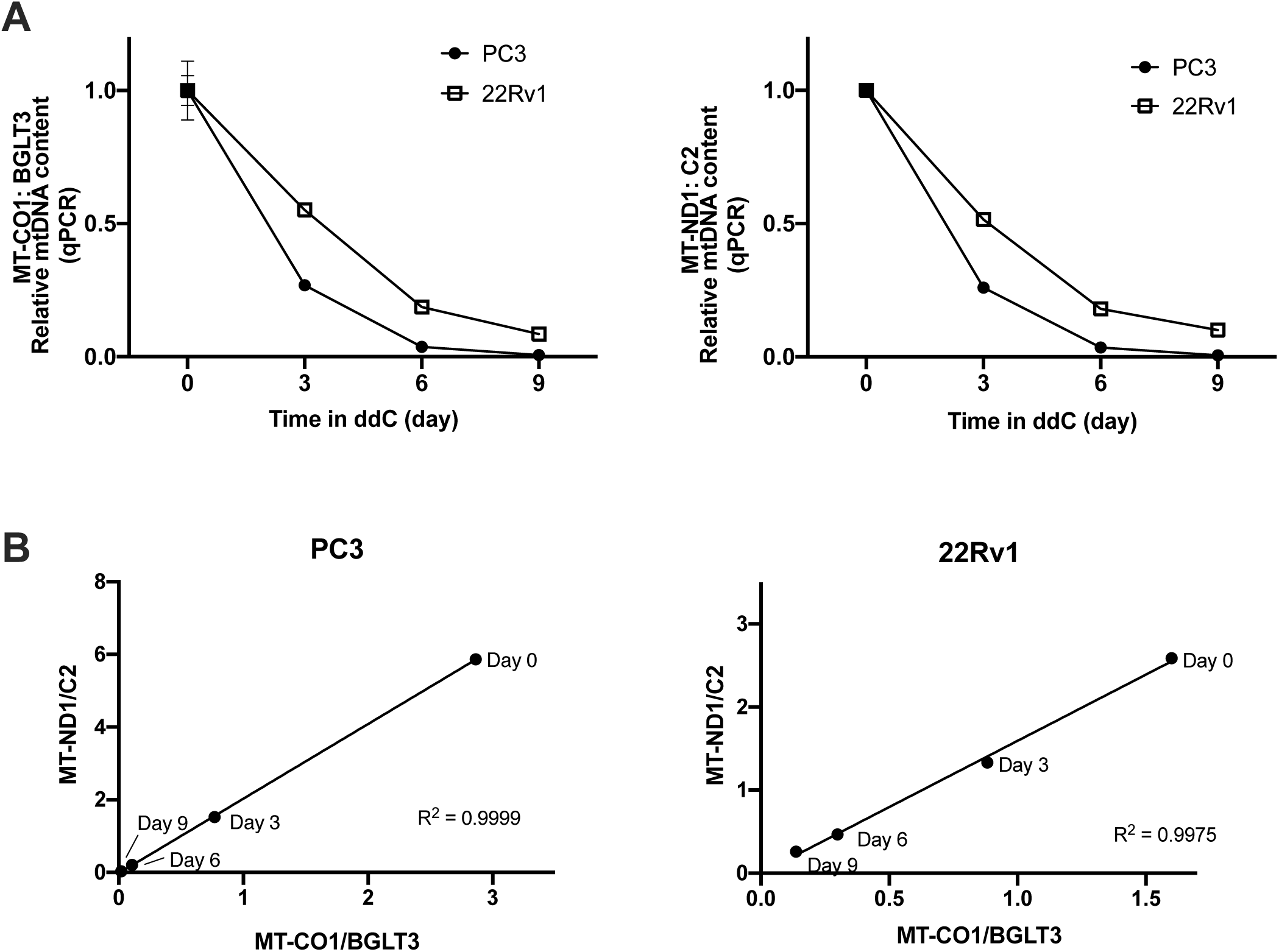
Time-dependent reduction in mtDNA copy number measured by quantitative PCR in prostate cancer cell lines treated with ddC. ddC concentration: 3 uM for PC3, 30 uM for 22Rv1. **A**: Stepwise reduction in the ratio of mtDNA to nuclear DNA for two different primer/probe sets. **B**: The ratios of mtDNA/nuclear DNA content from these two sets of primer pairs were highly correlated. Relative standard curve method was used to calculate the quantity of each amplified product. Fold differences were determined by defining the ratio of mtDNA gene to nuclear DNA gene in day 0 cells as 1.0. All qPCR reactions including the standard curve were performed in triplicate. Data are represented as means ± SD.

**Figure 4.**
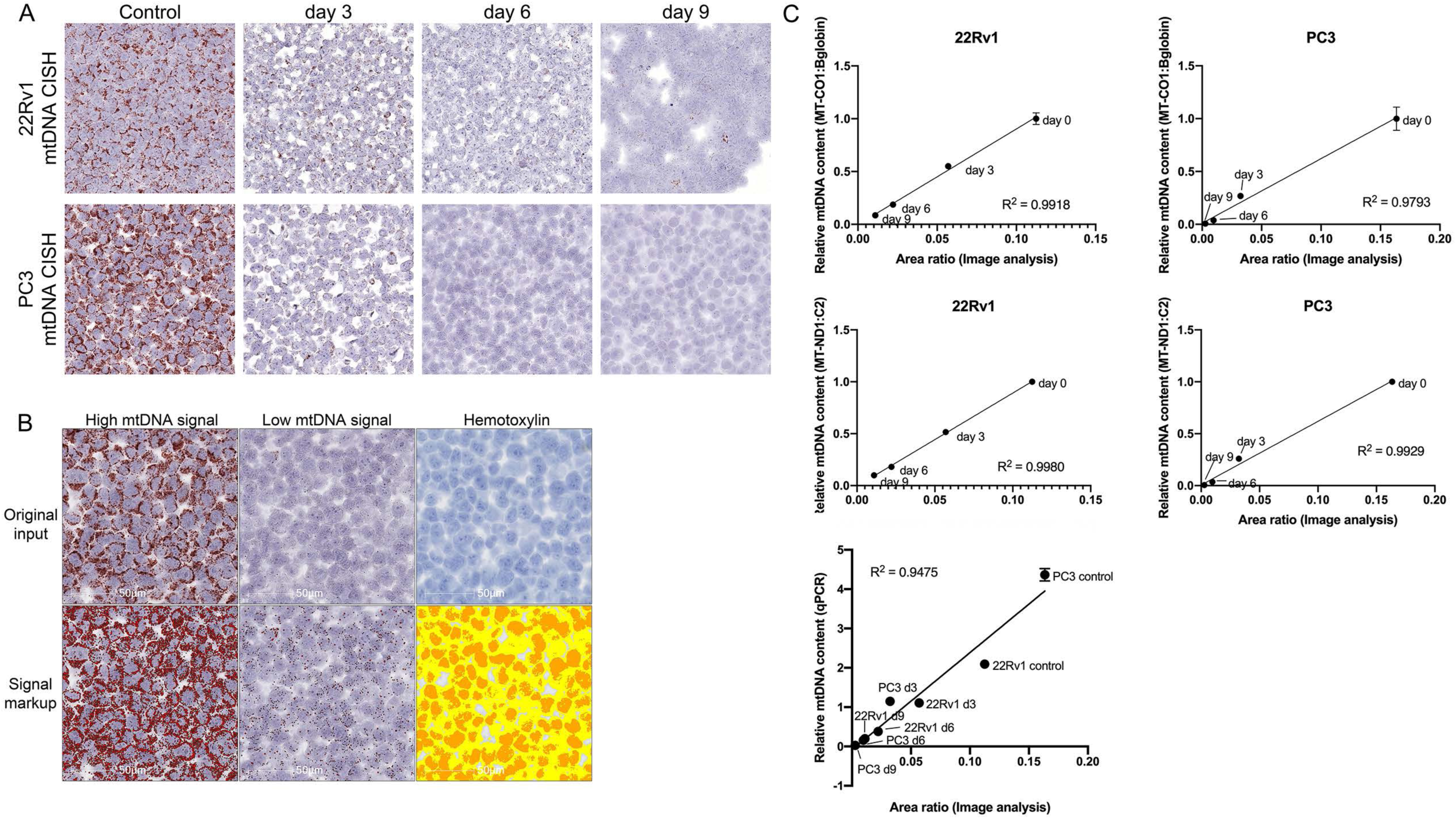
Quantitative nature of the *in situ* assay for mtDNA copy number determined by strong positive correlation between qPCR and the area-fraction of mtDNA signals by image analysis. **A**: Stepwise decrease in mtDNA *in situ* signals in relation to duration of drug treatment in FFPE cell blocks. **B**: Example of Halo™ interface for image analysis. **C**: Correlations between relative mtDNA content by qPCR and by area-fraction using Halo^TM^.

When mtDNA is depleted, the mtDNA encoded mRNAs and proteins will also be greatly reduced.^27, 46^ As such, we found that the loss of mtDNA was accompanied by a marked reduction of the mtDNA encoded protein MT-CO1 in a time-dependent manner in response to ddC treatment, whereas the nuclear DNA-encoded COX-IV protein remained present as expected (**Supplementary Figure 1**).

The sequence identity between human mtDNA and mouse mtDNA in the targeted regions of the *MT-CO1* gene is 77.8%. A prior study using RNAscope technology to distinguish mouse and human gene expression at the RNA level showed that for probe sets with <85% homology the probes did not cross hybridize.^47^ Therefore, we further tested the specificity of the hybridization using human cancer cells grown as xenografts in mice. Hybridization signals for the human *MT-CO1* probe set were observed exclusively in the human tumor cells but not in the mouse stromal cells (**Supplementary Figure 2**). Conversely, a probe set developed to the mouse *mt-Co1* gene showed signals only in the mouse stromal cells and not in the human cells (**Supplementary Figure 2**). This result is consistent with the high specificity of these DNAscope hybridization reactions.^37, 47, 48^

To verify whether the *in situ* signals were localized to mitochondria, we used a fluorescence approach for *in situ* hybridization combined with immunofluorescence against MRPL12, a well-known mitochondrial protein.^39, 40, 49^ As visualized by confocal microscopy (**Supplementary Figure 3**), the punctate signals for mtDNA and MRPL12 protein showed prominent areas of overlap in the cytoplasm, consistent with mitochondrial localization.

### mtDNA Copy Number Survey of Normal Tissues by in situ Hybridization

Cellular and tissue heterogeneity of mtDNA copy number is poorly explored. We applied mtDNA CISH to various tissues from mice and humans. As an example of marked tissue/cell type heterogeneity, Figure 5 shows a cross section from the neck region of a mouse encompassing a range of normal tissues. Strong mtDNA signals were present in the basal layer (Figure 5 C-D and see below regarding stem cell compartments) of squamous epithelium in the esophagus, as well as in skeletal muscle, and glandular tissues including the tracheal epithelium, submucosal glands and thyroid (Figure 5). Very low hybridization signals were seen in tracheal cartilage (Figure 5B), consistent with its known lack of internal blood vessels and low mitochondrial mass. Brown adipose tissue (Figure 5 E-F) showed very high levels of mtDNA signals, in line with the well-known high levels of mitochondria in this tissue, compared to white adipose tissue.^50–52^ Salivary glands showed a pattern in which the ducts showed higher mtDNA signals compared to the acini (not shown).

**Figure 5.**
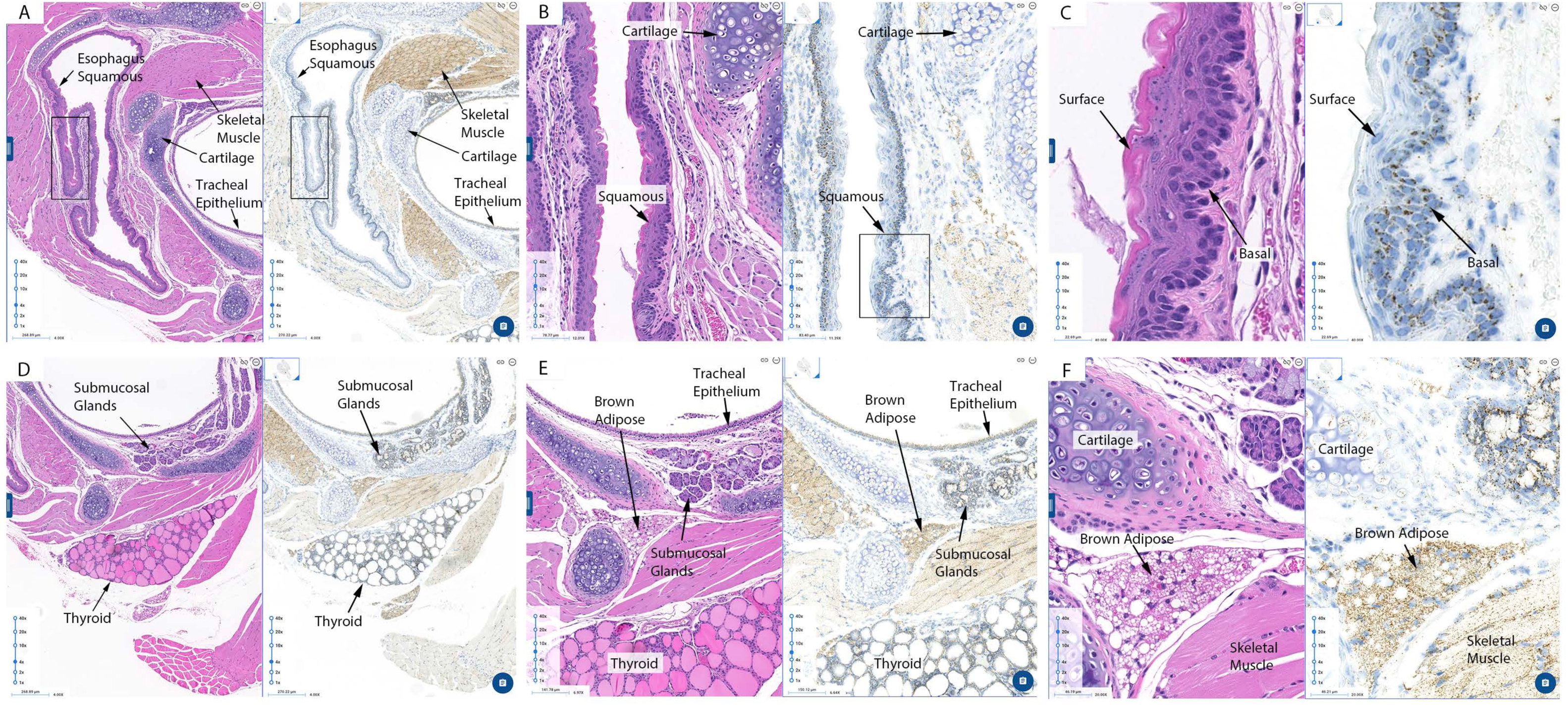
Cross section from the neck region from a 2-month-old female FVB/N female mouse. **A-F**: Strong mtDNA signals were present in the basal compartment of squamous epithelium in the esophagus, as well as in skeletal muscle, and glandular tissues including the tracheal epithelium, submucosal glands and thyroid. Very low hybridization signals were seen in tracheal cartilage (**B**). Brown adipose tissue also showed very high levels of mtDNA signals (**E-F**).

As expected, we found relatively high levels of mtDNA hybridization signals in cardiac muscle tissue, commensurate with their high energy demands and known high mitochondrial mass (not shown). In the pancreas, the islets showed more mtDNA signals than the exocrine acini (not shown). This corresponds to the observations of higher mitochondrial protein expression in pancreatic islets versus acini.^53^ In the urinary bladder, there were strikingly high levels of mtDNA signals in umbrella cells in female mice, as compared to the underlying intermediate or basal cells, which were higher than the detrusor muscle and other mesenchymal tissues (**Supplemental Figure 4**).

### Adrenal Gland

The mouse adrenal showed marked differences in mtDNA signals by zone. In the cortex there appeared to be a slight gradation of signals, with somewhat stronger signals towards the capsule in the zona glomerulosa, and a tapering of signals in the zona fasciculata towards the medulla (Figure 6). Mice do not have a zona reticularis,^54^ but in female mice the X-zone showed higher levels of mtDNA than the zona fasciculata. Strikingly, the adrenal medulla showed much lower mtDNA signals than the cortex (Figure 6), in both males and females, which is consistent with prior results by qPCR.^55^

**Figure 6.**
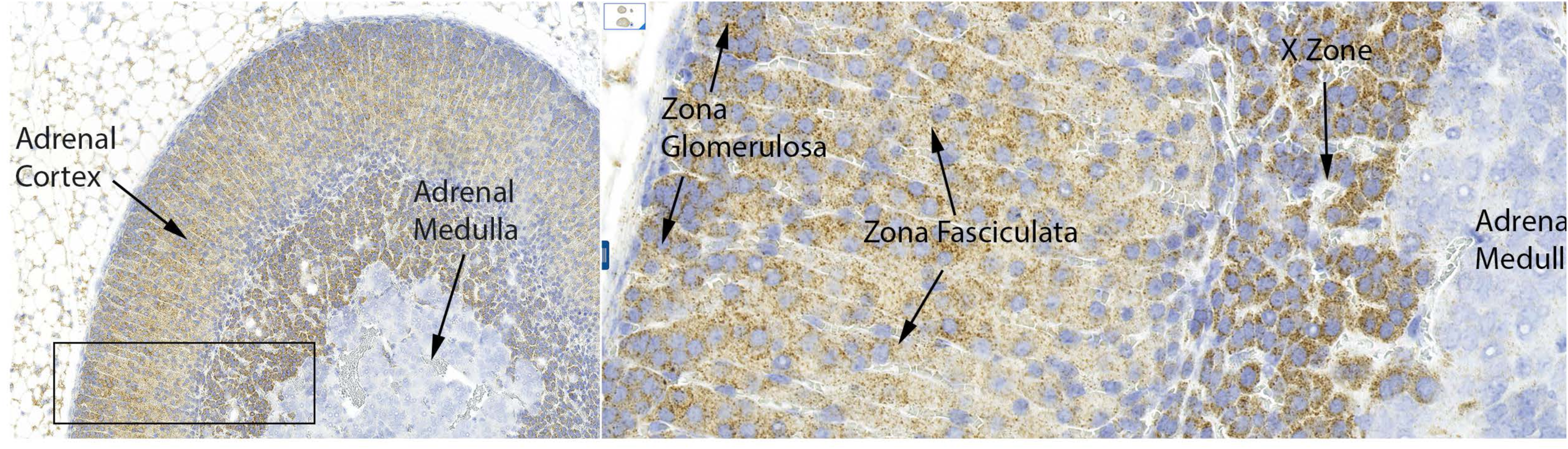
Cross section of an adrenal gland from a 2-month-old female FVB/N mouse. The adrenal medulla showed much lower mtDNA signals than the cortex. The cortex showed variation in different zones with prominent *in situ* signal in the X-zone. Original magnification x10 (left), x40 (right).

### Kidney

In the mouse kidney we found robust signals in cortical tubular epithelial cells and strikingly low signal levels in cells within glomeruli (Figure 7). Distal convoluted tubular epithelium showed stronger signals than in proximal tubules. In collecting ducts, present in the cortex in medullary rays, and in the medulla, there were markedly reduced mtDNA hybridization signals compared to proximal and distal tubules. We found a highly similar pattern in human kidney tissues (not shown). As an additional control for hybridization efficiency in the different renal parenchymal compartments, we performed DNA *in situ* hybridization for telomeres using a highly similar chromogenic approach.^34, 56^ We found similarly strong telomeric DNA signals in glomeruli as well as throughout the different renal tubules, supporting the likelihood that lower levels of mtDNA signals are not related to failure of probe penetration (not shown).

**Figure 7.**
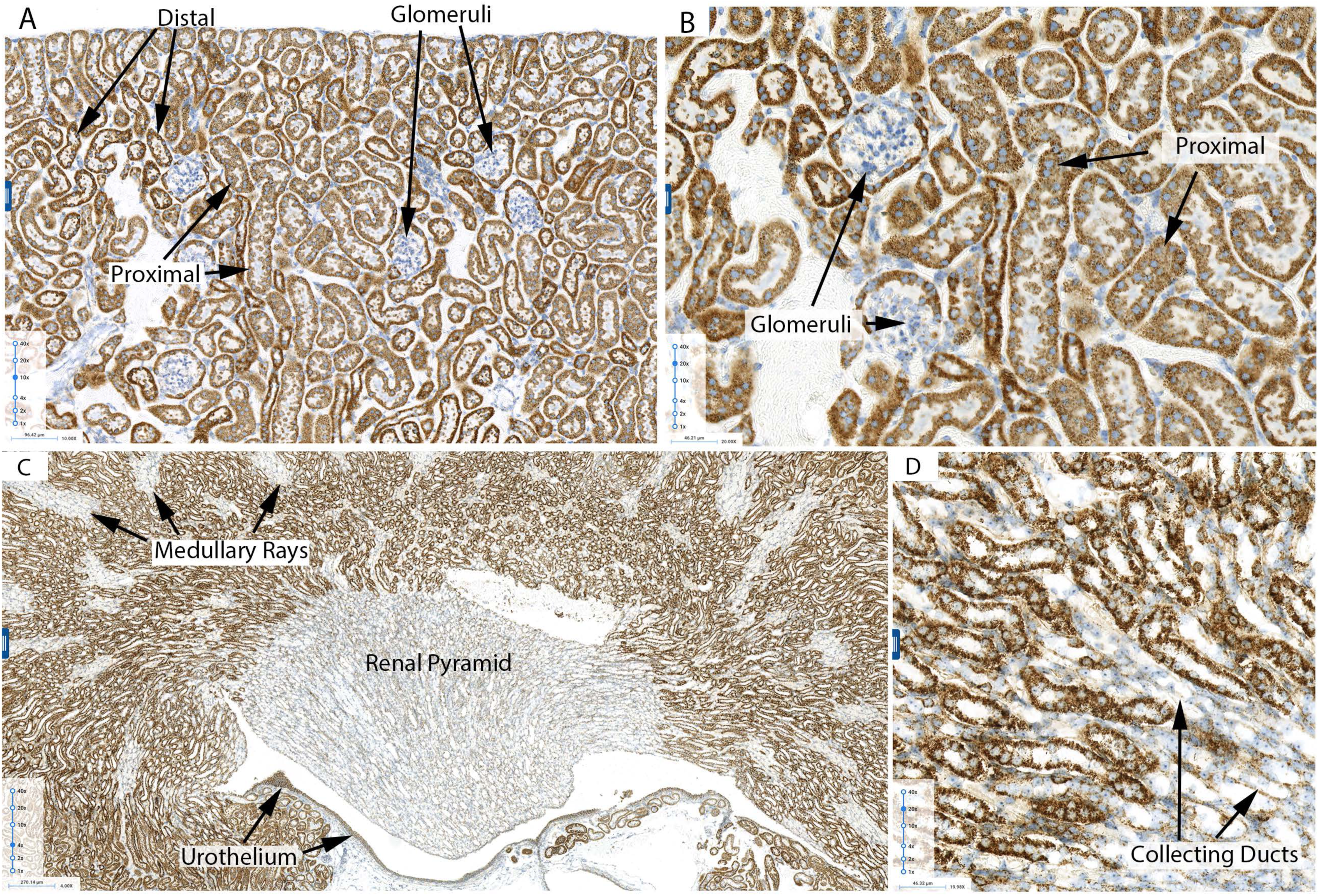
Cross section of a kidney from a 2-month-old male FVB/N mouse. **A-B**: Very low hybridization signal levels were present in the glomeruli. Epithelial cells in distal convoluted tubules showed higher mtDNA signals than those in proximal tubules. **C-D**: Collecting ducts in medullary rays and in the medulla showed notably reduced mtDNA hybridization signals.

### Liver

In the mouse liver there was a striking zonal distribution to mtDNA hybridization signals such that periportal hepatocytes (zone 1) showed the highest levels, with an apparent gradient of lower levels towards the hepatocytes surrounding the central veins (zone 3) (Figure 8). This result is consistent with a recent study by Brosch et al. who examined next generation DNA sequencing data for mtDNA after laser capture microdissection of different human liver zones.^21^

**Figure 8.**
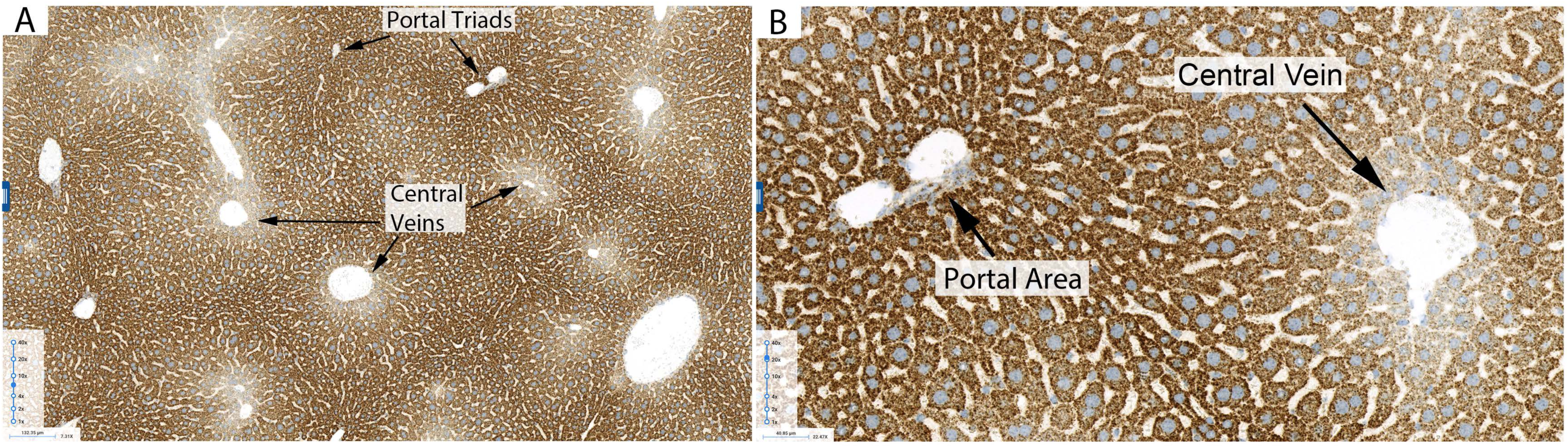
Cross section of liver from a 2-month-old male FVB/N mouse. There was a zonal distribution to mtDNA hybridization signals such that higher levels were present in the periportal hepatocytes and lower levels in the pericentral ones (**A**). **B** shows a higher magnification image of **A**.

### Self-renewing Tissues with Stem Cell Compartmentalization

Based on the findings above in the esophagus (Figure 5 **above**), we performed mtDNA CISH in other continuously self-renewing tissues containing hierarchical differentiation, starting with other stratified squamous epithelia. In each stratified squamous tissue examined, including mouse skin, cervix, vagina, oropharynx, and esophagus, the cells in the basal compartment showed higher mtDNA signals than cells toward the surface (**Supplemental Figure 5** shows cervix; other sites not shown). In mouse hair follicles, those that are in the anagen growth phase showed intense signals in the deepest regions with a gradation of signals towards the skin surface (**Supplemental Figure 6)**. Also, these anagen phase regions showed considerably higher mtDNA signals compared with those in telogen phase (**Supplemental Figure 6)**. This result agrees with previous studies showing higher mitochondrial activity by Mitotracker in anagen cells than telogen cells.^57^ In the gastrointestinal tract, the human (Figure 9) and mouse small intestine showed more mtDNA signals in the crypts, where the stem cells and transient amplifying/proliferative compartment cells are located, compared to the villus tips (Figure 9). A similar pattern was present in the colon of both mouse and human, and mouse stomach, where the stem/proliferative cells in the isthmus showed higher mtDNA signals than the surface or deeper glands (**Supplemental Figure 7).** We conclude that in each rapidly self-renewing tissue type that is organized using hierarchical differentiation, the stem cell/proliferative cell compartments contain higher mtDNA signals compared to the more differentiated cells.

**Figure 9.**
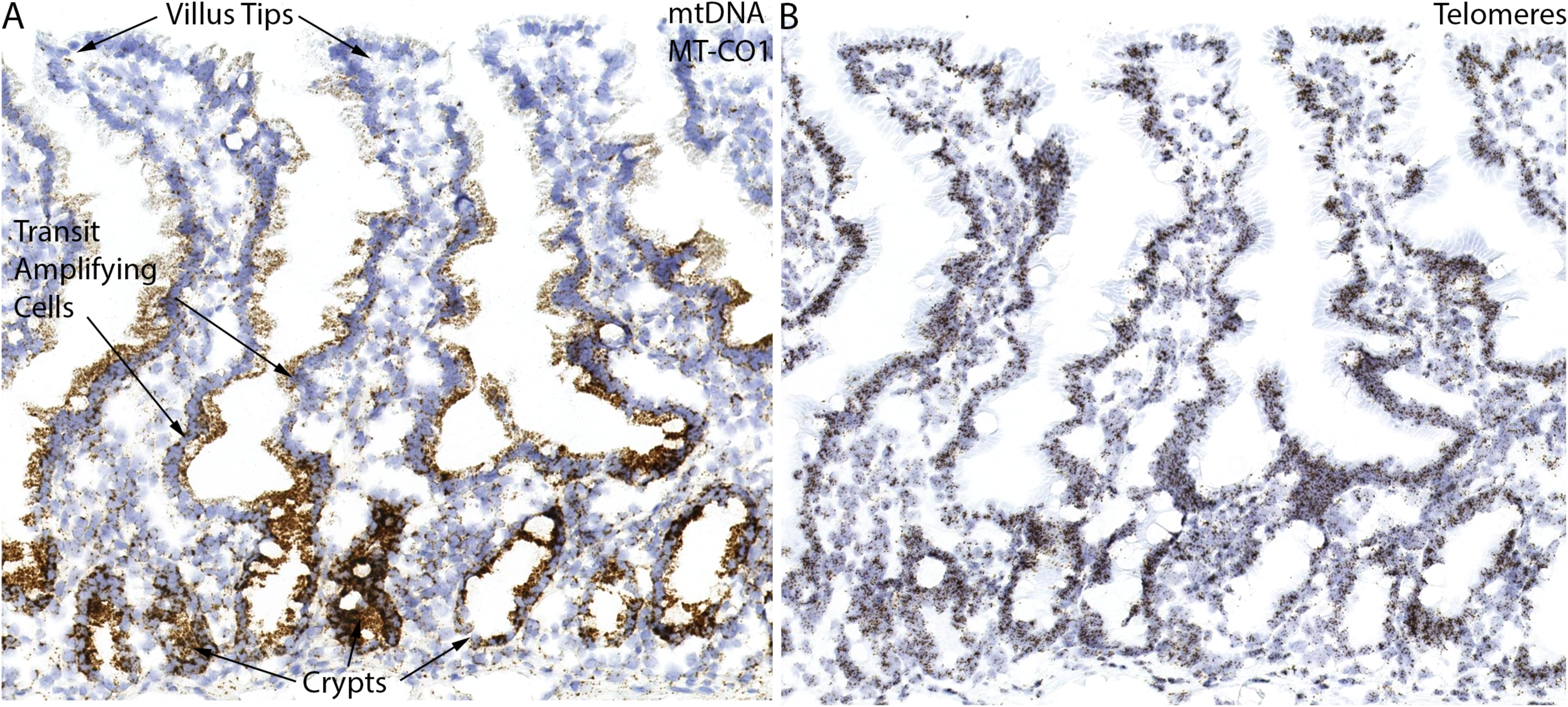
*In situ* hybridization staining of mtDNA and telomeres on frozen human duodenum. **A**: Epithelial cells in the crypts and transit/amplifying cell compartments had higher mtDNA *in situ* signals than to those in the villus tips. **B**: Telomeric DNA *in situ* hybridization on adjacent slides of **A** as a control for probe penetrance. Strong telomeric DNA signals were seen through the bottom of the crypts to the top of the villus. **C** and **D** are higher magnification images of **A** and **B**, respectively. Original magnification x10 (**A** and **B**), x20 (**C** and **D**).

### Male Reproductive Tissues

In the prostatic epithelium, basal cells contained increased signals in general as compared to luminal cells (**Supplemental Figure 8**), and the epithelium showed overall higher signals than the stroma. The higher mtDNA content in the basal cell layer might be due to the fact that MYC is almost exclusively expressed in the basal cells,^58–60^ and MYC has been shown to regulate mitochondrial biogenesis,^58^ raising the question that mtDNA levels in some tissues are physiologically regulated by MYC (see Discussion). In the seminiferous tubules of the testis, we observed strong signals in germ cells adjacent to the basement membrane with an overall decrease in signals as cells differentiated towards mature sperm in the lumens (**Supplemental Figure 9**), which is consistent with prior studies,^61, 62^ In the epididymis, there were more mtDNA signals in the tail epithelium as compared with epithelium in the head (**Supplemental Figure 9**).

### Female Reproductive and Urinary Tissues

The epithelium in mouse fallopian tube and uterus had strong mtDNA *in situ* signals (**Supplemental Figure 10**). In the mouse ovary, in regions containing follicles, there were very strong signals in the granulosa cells directly surrounding the oocytes, with many, albeit less numerous, signals in the oocyte cytoplasm. In addition, immature corpora lutea showed higher mtDNA signals compared to the mature corpora lutea (**Supplemental Figure 10**). Mouse epithelial cells in the endometrium showed clearly higher mtDNA signals as compared with endometrial stroma (**Supplemental Figure 10**).

### Central Nervous System

In the central nervous system, neuronal cell bodies generally contained much stronger signals than surrounding glial cells and white matter. **Supplemental Figure 11** illustrates this in the mouse cerebellum which shows strong signals in Purkinje cell neuronal cell bodies and in neuronal cell bodies within a deep cerebellar nucleus.

### Aging Kidney and Calorie Restriction

Kidney function is known to decrease gradually with increasing age in mammals, which has been associated with a gradual decline in mitochondrial function.^63–65^ A recent study found that the ratio of mtDNA to nuclear DNA decreased in the aging mouse kidney.^36^ Further, this reduction was prevented by calorie restriction, which is consistent with the finding that calorie restriction can prevent age-related declines in mitochondrial biogenesis and function in the kidney.^36^ However, which compartments in the kidney are most affected by these changes were not explored. Therefore, we examined the effects of aging and calorie restriction on mouse mtDNA levels in the kidney using our *in situ* assay. Mice were either fed *ad libitum* or treated with calorie restriction and then aged to 24 months. Tissue sections from the mouse kidneys were hybridized for mtDNA. After whole slide scanning and blinding of the slides, a visual microscopic review was conducted by two separate observers. Using this blind review, each observer separated the mice into two groups. One group of 3 mice contained variably lower levels of mtDNA in all tubule compartments by visual estimate (Figure 10 shows cortical tubules) and was composed of the aging mice fed *ad libitum*. Although a general reduction in mtDNA signals was present throughout the kidney, in some of the cortical tubules there were markedly reduced signals, and this was only seen in the *ad libitum* fed older mice (Figure 10). The other group (N=6), which did not show these isolated areas of markedly reduced mtDNA signals, consisted of a combination of younger mice fed *ad libitum* and aged calorie-restricted mice; these two treatment groups were visually indistinguishable.

**Figure 10.**
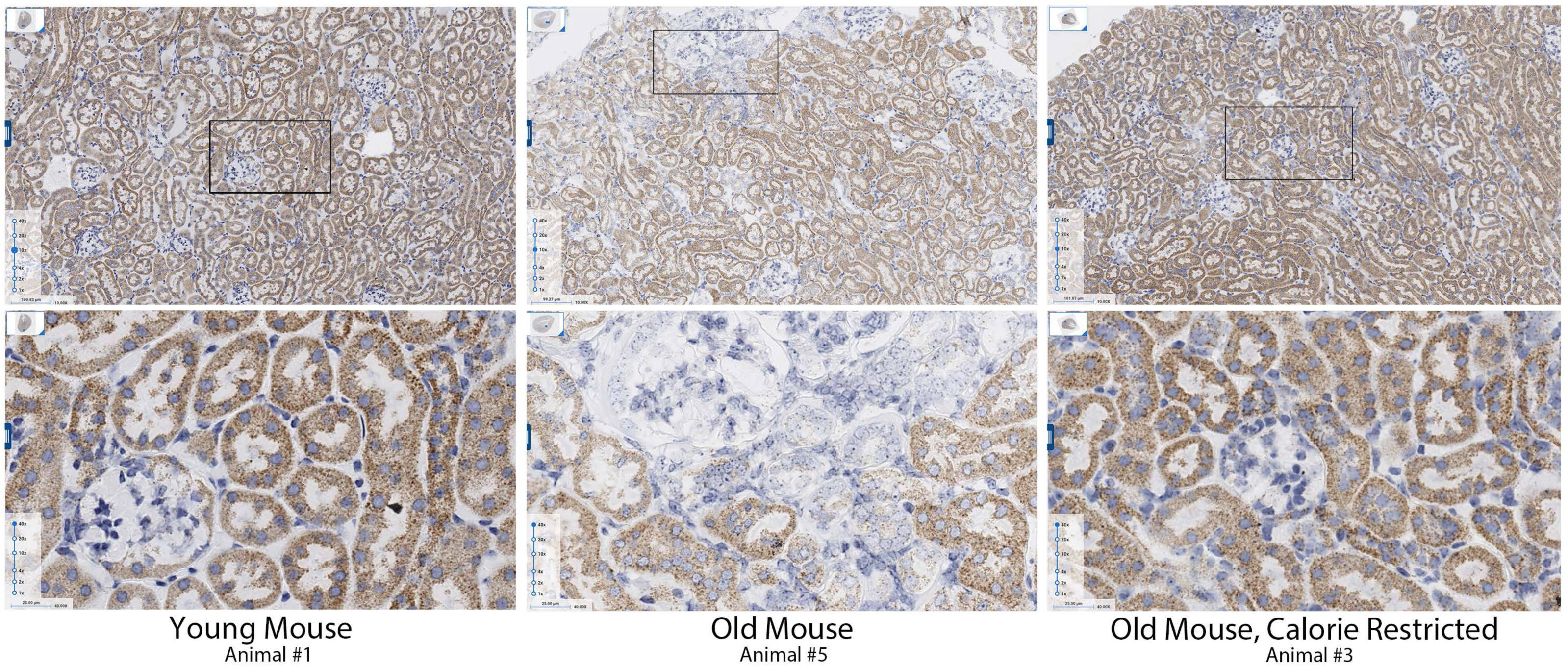
mtDNA *in situ* hybridization in kidneys from young, aging, and aging with calorie restriction C57BL/6 mice. In the aging mouse kidney, decreased mtDNA levels were identified in tubules, which was prevented by calorie restriction. n=3 for each group. Bottom panel is higher magnification images of top panel. Original magnification x10 (top panel), x40 (bottom panel).

## DISCUSSION

There is no gold standard for measuring cell-type heterogeneity in mtDNA copy number in tissue samples. We present a novel approach that is widely applicable, straight-forward to implement and can be used either qualitatively or quantitatively. A key finding using this approach, that was not previously appreciated, is that in tissues organized into self-renewing hierarchical stem cell driven differentiation systems,^66^ cells within stem cell/proliferative compartments contain higher mtDNA levels compared to their more differentiated progeny. This finding raises a number of important questions. The first involves our understanding of mtDNA replication. While it is known that mtDNA replication is uncoupled from cell cycle progression (e.g. nuclear DNA replication) in some mitotic and post mitotic cells,^67, 68^ a number of groups have shown mtDNA replication appears to be synchronized during the cell cycle in cell culture systems.^31–33, 69^ Thus, the findings to date are somewhat contradictory. Future studies using an *in situ* approach, coupled with the ability to visualize newly synthesized mtDNA and nuclear DNA in intact tissues, can help clarify this question.

Another question is how mtDNA levels are regulated in the self-renewing tissues described herein. Key regulators of mtDNA replication are peroxisome proliferator activated receptor coactivator 1 alpha (PGC1-ɑ; encoded by *PPARGC1A*) and peroxisome proliferator activated receptor coactivator 1 beta (PGC-1β; encoded by *PPARGC1B*). Further, MYC has been shown to drive mtDNA replication and mitochondrial biogenesis.^58^ Interestingly, in the intestine^70^ and squamous epithelia,^71^ MYC levels are highest in regions that clearly overlap with those we have found have the highest mtDNA levels, raising the possibility that physiological regulation of mtDNA during tissue renewal in many stem cell compartments is governed, at least in part, by MYC. In fact, in mice with targeted disruption of *Myc* in the intestine, Myc-deficient crypts are lost, and Myc is required for successful progression through the cell cycle.^70^ It is possible, therefore, that one mechanism by which Myc functions in the intestine is to induce mtDNA replication and mitochondrial biogenesis. Future studies are needed to address the question regarding the regulation of mtDNA levels in normal self-renewing tissues by MYC. Other genes related to mitochondrial function also appear to be required for intestinal stem cell maintenance since mice with targeted disruption of the key mitochondrial protein chaperone, hsp60, show loss of stem cells and proliferative capacity.^72^ Another question is why prior studies have not shown the stark differences we observed in stem/proliferative compartments in mtDNA levels.^72^ To gain some insight into this we are currently performing IHC against a number of mitochondrial enzymes and *in situ* hybridization for mtRNAs. Our preliminary findings in the human intestine is that that mitochondrial protein markers do not correlate with mtDNA levels, but that 12S rRNA (*MT-RNR1*) does.

In the kidney of young healthy mice and humans, striking differences in mtDNA levels within different renal epithelial compartments were observed. First, we verified much lower levels of mtDNA in virtually all cell types present within the glomeruli.^28^ Interestingly, distal tubules showed higher mtDNA levels than proximal, and collecting ducts showed markedly lower levels than proximal or distal tubules. Prior studies have shown a reduction in mtDNA to nuclear DNA by PCR based assays in aging mouse kidney tissues, and this could be prevented by calorie restriction,^36^ treatment with a dual agonist of the farnesoid X receptor (FXR) and the G protein-coupled receptor TGR4, INT-767,^36^ or the estrogen related receptors ERRɑ, ERRβ, and ERR*γ*.^65^ Consistent with this, in the present study we found that the marked reduction in mtDNA levels was visualized in a number of tubule compartments and was prevented by calorie restriction. The current *in situ* approach will be useful to determine whether similar changes are seen in the human kidney with aging and how mtDNA levels are altered in different renal pathologies.

Another interesting finding was the remarkable zonal differences in mtDNA numbers in hepatocytes that is quite consistent with the known oxygen gradient, as well as, a recent study using NGS from laser capture microdissected zones of the liver.^21^

The ability to routinely interrogate mtDNA levels at the cellular level in the context of normal and abnormal spatial organization of cell types should greatly facilitate approaches to deciphering changes that occur in mtDNA in a number of disease states, including different cancers,^73^ their precursor lesions and a large spectrum of non-neoplastic clinical disorders. These include inherited autosomal and mtDNA syndromes that affect mtDNA copy number^74^ that involve a number of organ systems and cell types (e.g. involving skeletal and cardiac muscle, neurons in Parkinson’s disease and other neurodegenerative disorders, hepatocytes in liver, etc.). We submit that *in situ* studies of the involved tissues using the current approach can facilitate novel insights into disease pathogenesis and response to therapies.

In summary, we present a novel, quantitative and straightforward approach to mtDNA level measurements in tissue samples. This approach provides the first widely available opportunity to test changes that occur in mtDNA copy number in specific cell types, while retaining spatial information. The current study uncovered evidence of increased mtDNA copy number in stem/proliferative compartments throughout the body and serves as an initial phase towards the completion of a comprehensive cellular atlas of mtDNA copy number across tissues, and changes in such cell types in aging and disease. It should complement approaches using genomic methods, such as those using single cell RNA-seq and ATAC-Seq in which several large-scale efforts are underway. It can also complement other *in situ* approaches such as spatial transcriptomics, highplex IHC and *in situ* hybridization and prove useful in studies related to the effects of genotype, environmental exposures, and interventions (e.g. dietary or exercise) in model systems and humans.

## ACKNOWLEDGEMENTS

This study was supported by the NIH/NCI SPORE in Prostate Cancer: P50CA58236, and the NIH/NCI U01 CA196390 for the Molecular and Cellular Characterization of Screen Detected Lesions (MCL), the U.S. Department of Defense Prostate Cancer Research Program (PCRP), The Johns Hopkins Sidney Kimmel Comprehensive Cancer Center Oncology Tissue Services Laboratory supported by NIH/NCI grant P30 CA006973.

## SUPPLEMENTAL FIGURE LEGENDS

**Figure S1.** A reduction of the mtDNA encoded protein MT-CO1 accompanied the loss of mtDNA in a time-dependent manner in response to ddC treatment. The COX-IV protein remained present. Original magnification x40.

**Figure S2.** In human cancer cell xenografts, the mouse mtDNA probes only bound to the mouse stromal cells, and the human mtDNA probes only bound to human tumor cells.

**Figure S3.** Colocalization of MT-CO1 *in situ* signals (mtDNA) and MRPL12 protein. Note the puncta nature of mtDNA signals present exclusively in the cytoplasm.

**Figure S4.** Representative staining of mtDNA in the bladder of a 2-month-old female FVB/N mouse. High levels of mtDNA signals were present in umbrella cells compared to the underlying intermediate or basal epithelial cells. Original magnification x4 (left), x40 (right).

**Figure S5.** Representative staining of mtDNA in the normal uterine cervix. The cells in the basal compartment showed higher mtDNA signals than cells toward the surface.

**Figure S6.** Representative staining of mtDNA in different stages of hair growth cycle of mouse. Hair follicles in anagen growth phase (**A** and **B**) showed intense signals in the deepest regions with a gradation of signals towards the skin surface. Hair follicles in telogen phase had considerably lower mtDNA *in situ* signals (**C**). Mammary glands (inset in **C**) also stained with strong mtDNA *in situ* signals, similar to other glandular tissues. Original magnification x4 (**A** and **C**), x20 (**B**).

**Figure S7.** Representative staining of mtDNA in stomach from a 3-month-old male FVB/N mouse. The stem/proliferative cells in the isthmus showed higher mtDNA signals than the surface or deeper glands.

**Figure S8.** Representative staining of mtDNA in the normal human prostate. Basal cells contained increased mtDNA signals as compared to luminal cells. The stromal cells showed overall lower signals than the prostate epithelium.

**Figure S9.** Representative staining of mtDNA in the mouse male reproductive system. In the seminiferous tubules of the testis, strong signals were present in germ cells adjacent to the basement membrane, and signals decreased as cells differentiated towards mature sperm in the lumens. In the epididymis, there were more mtDNA signals in the tail epithelium than those in the head.

**Figure S10.** Representative staining of mtDNA in the mouse female reproductive and urinary system. The epithelium in mouse fallopian tube and uterus endometrium had strong mtDNA *in situ* signals. In the mouse ovary, in regions containing follicles, there were very strong signals in the granulosa cells directly surrounding the oocytes, with many, albeit less numerous, signals in the oocyte cytoplasm. Immature corpora lutea showed higher mtDNA signals compared to the mature corpora lutea.

**Figure S11.** Representative staining of mtDNA in the mouse cerebellum. Strong signals were observed in purkinje cell neuronal cell bodies and in neuronal cell bodies within a deep cerebellar nucleus.

