## Supplemental Figures for "An *in situ* atlas of mitochondrial DNA in mammalian tissues reveals high content in stem/progenitor cells"

**Figure S1**

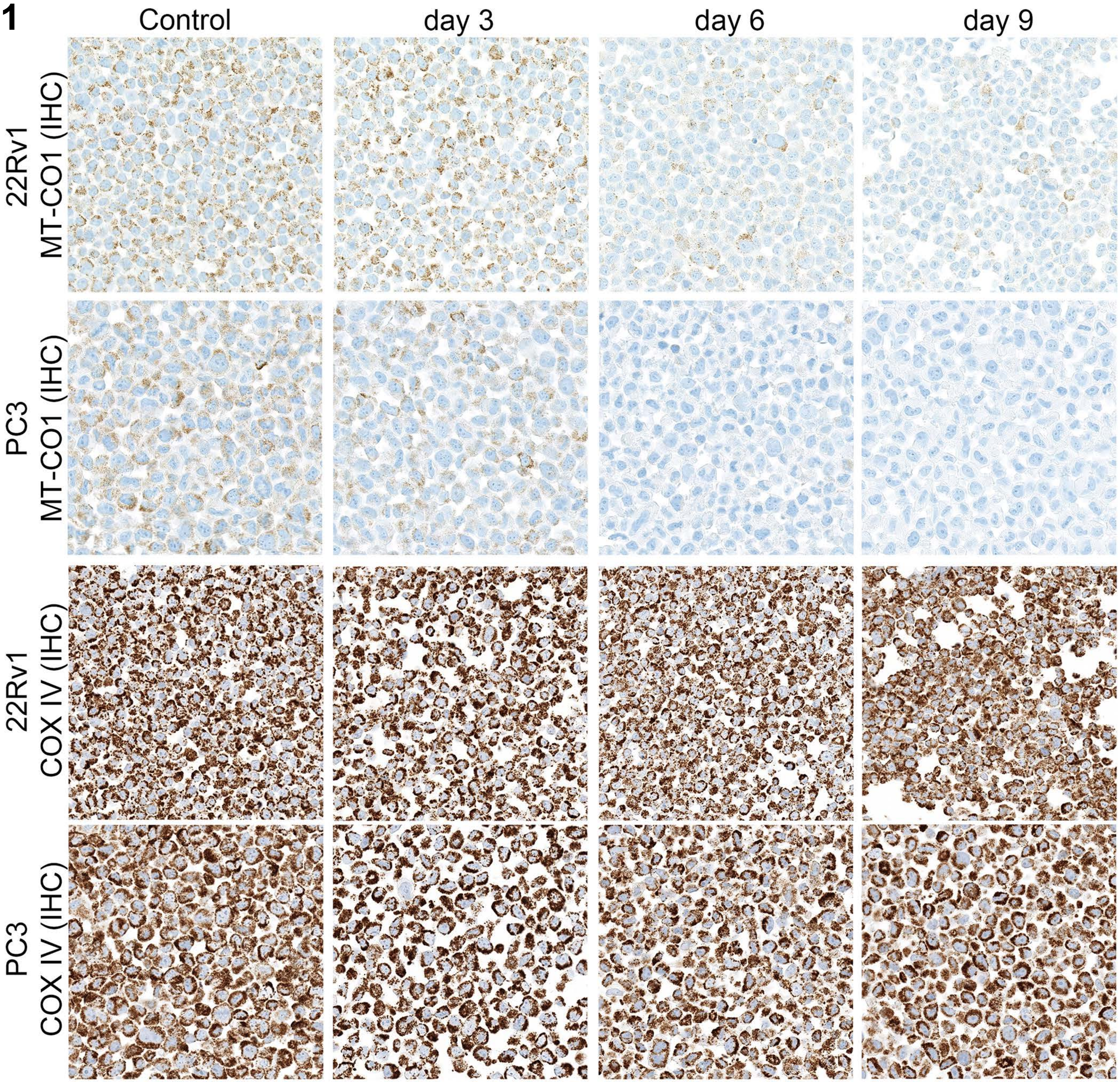
Control
day 3
day 6
day 9
22Rv1
MT-CO1 (IHC)
PC3
MT-CO1 (IHC)
22Rv1
COX IV (IHC)
PC3
COX IV (IHC)

#### Figure S2

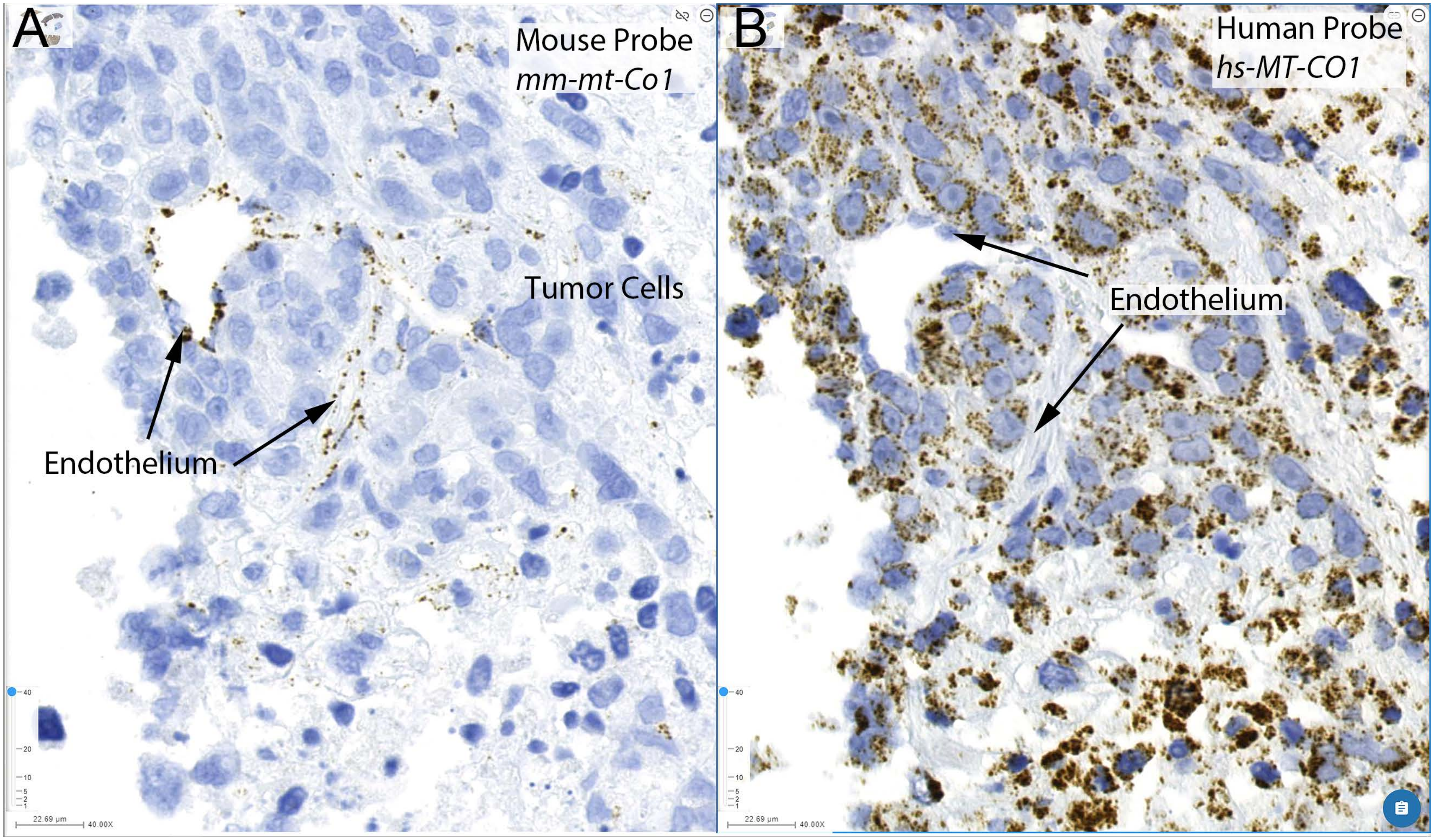

A
Mouse Probe
mm-mt-Co1
Tumor Cells
Endothelium
B
Human Probe
hs-MT-CO1
Endothelium

**Figure S3**

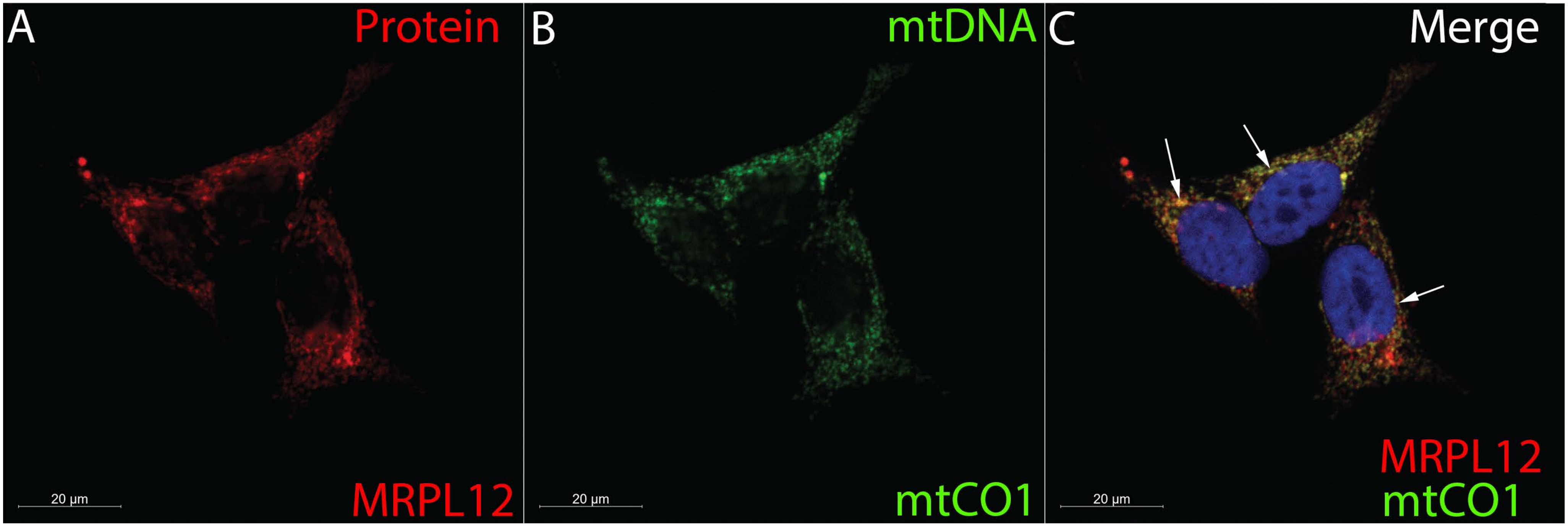
A
Protein
MRPL12
20 µm
B
mtDNA
mtCO1
20 µm
C
Merge
MRPL12
mtCO1
20 µm

#### Figure S4

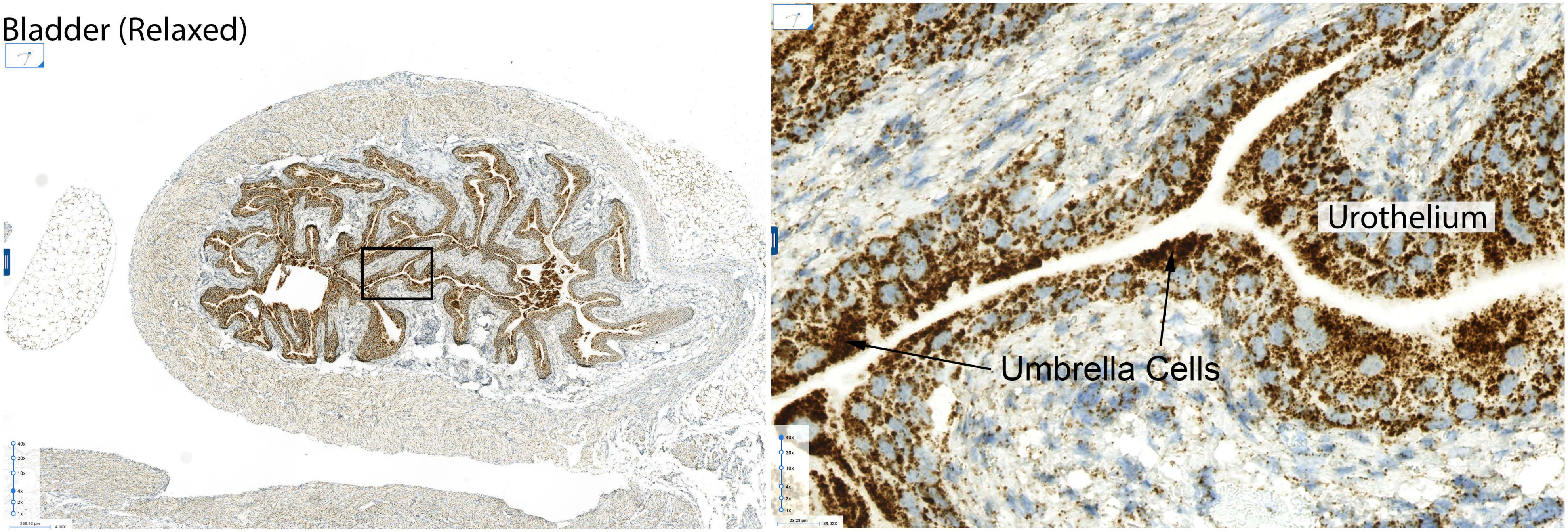

### Figure S5

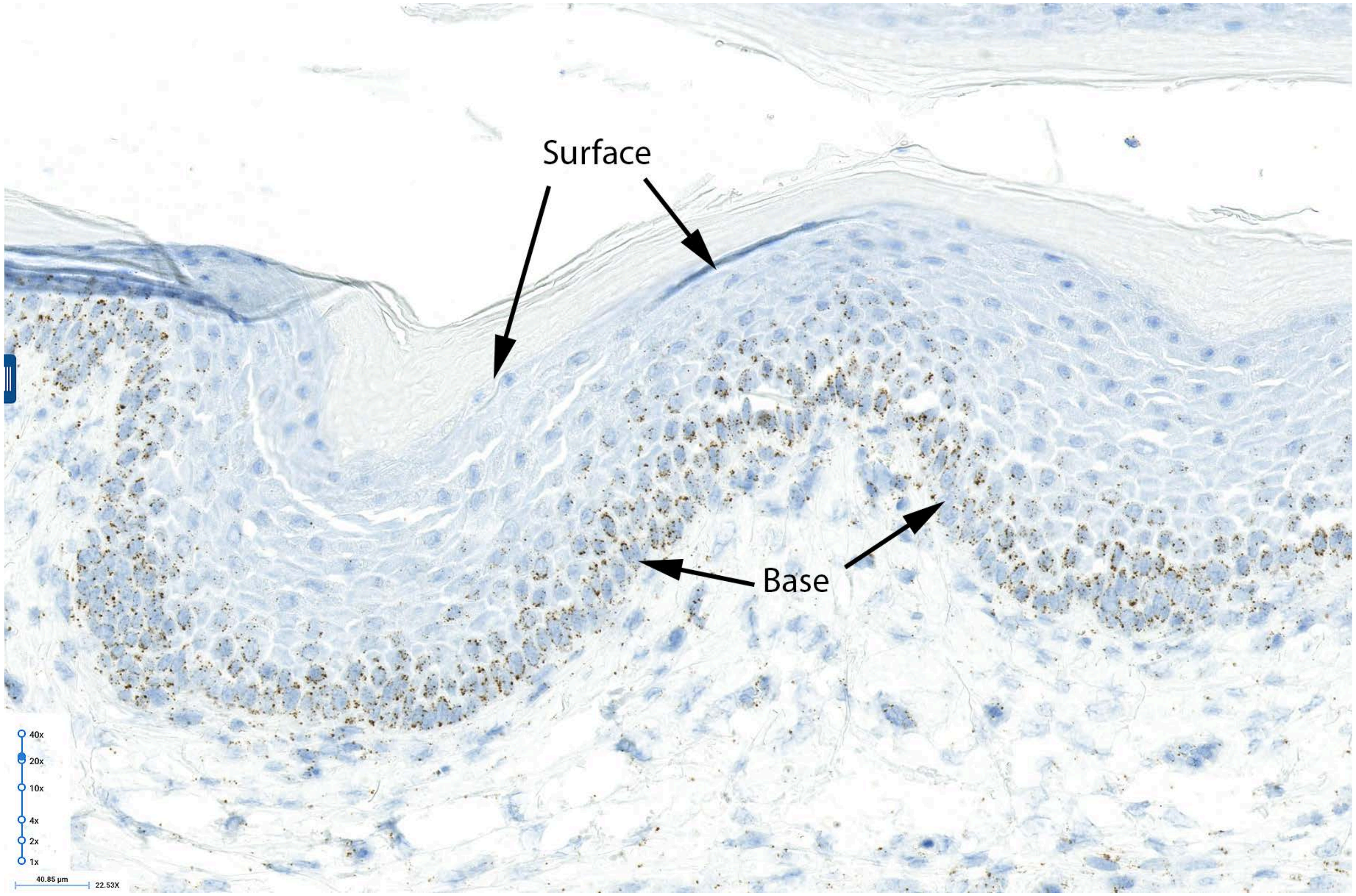

### Figure S6

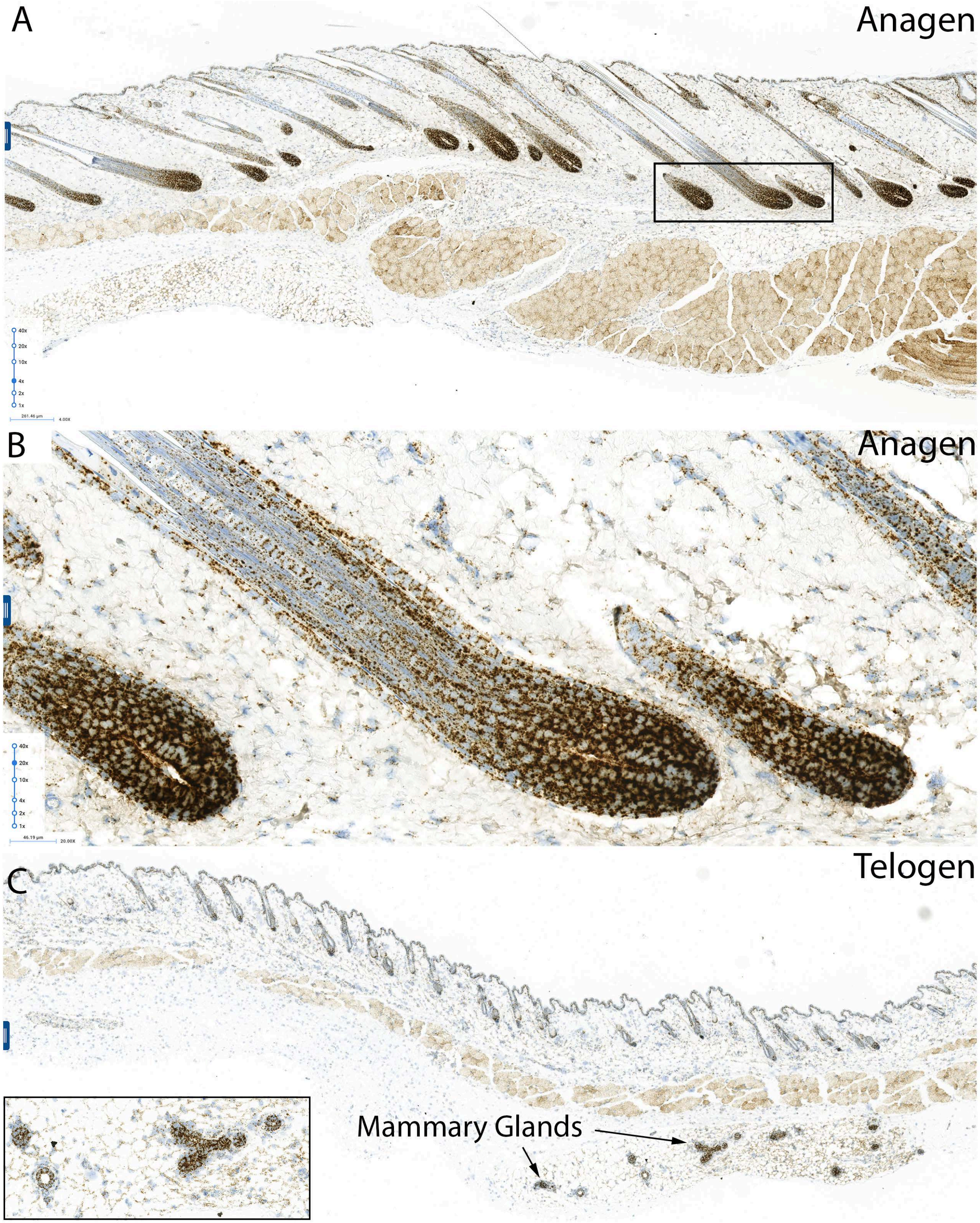

**Figure S7**

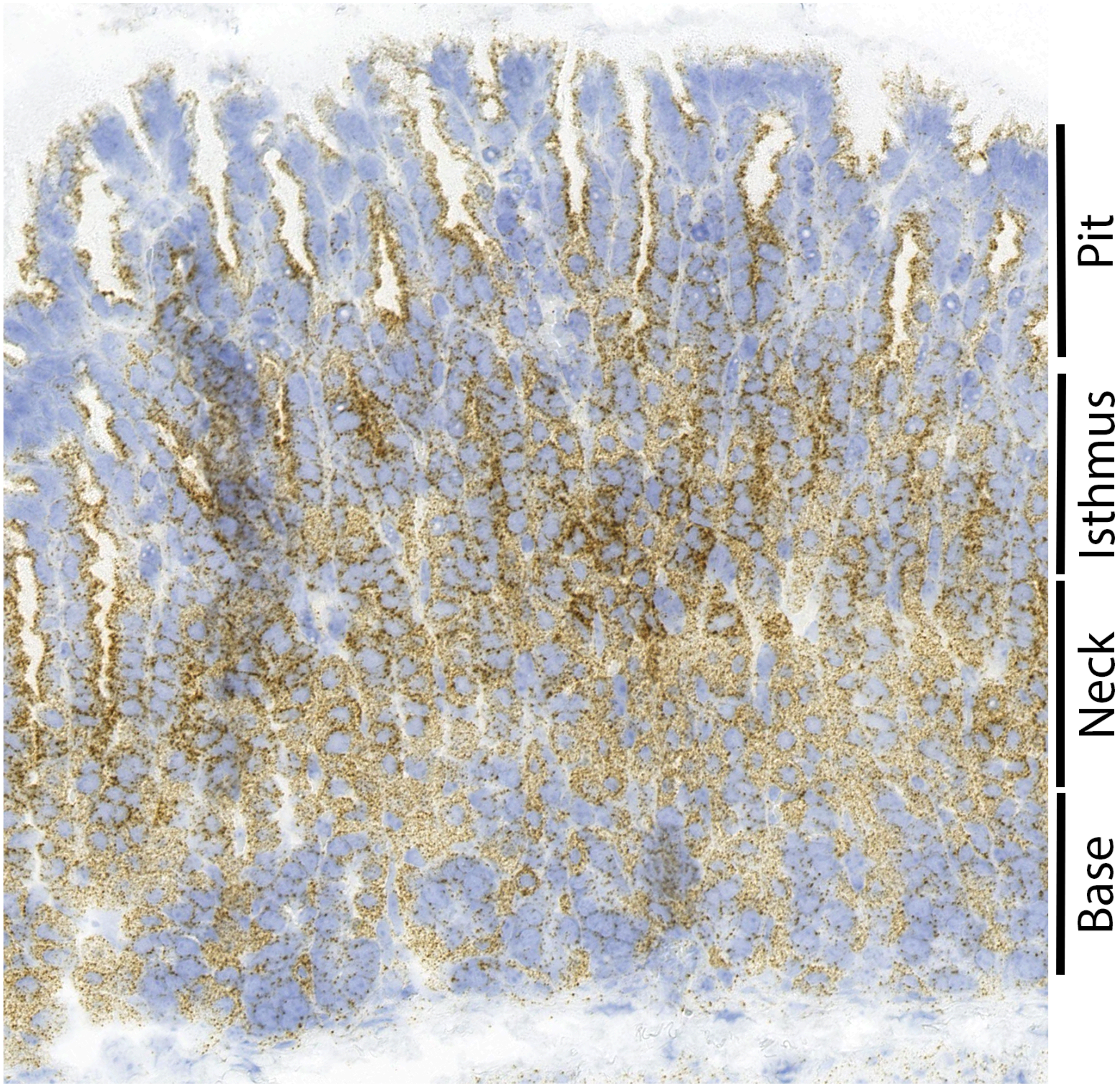

### Figure S8

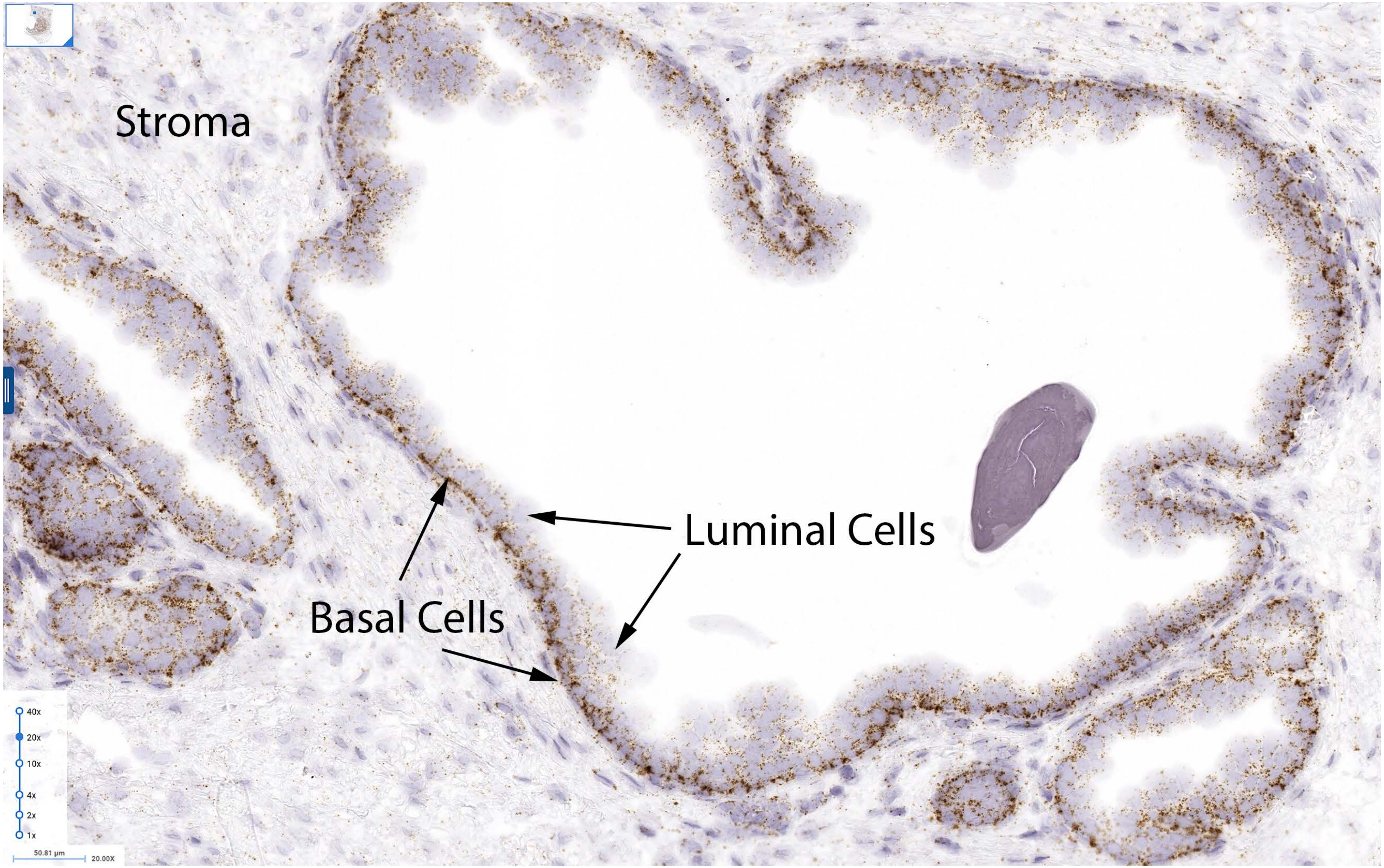
Stroma
Luminal Cells
Basal Cells
40x
20x
10x
4x
2x
1x
50.81 µm
20.00X

#### Figure S9

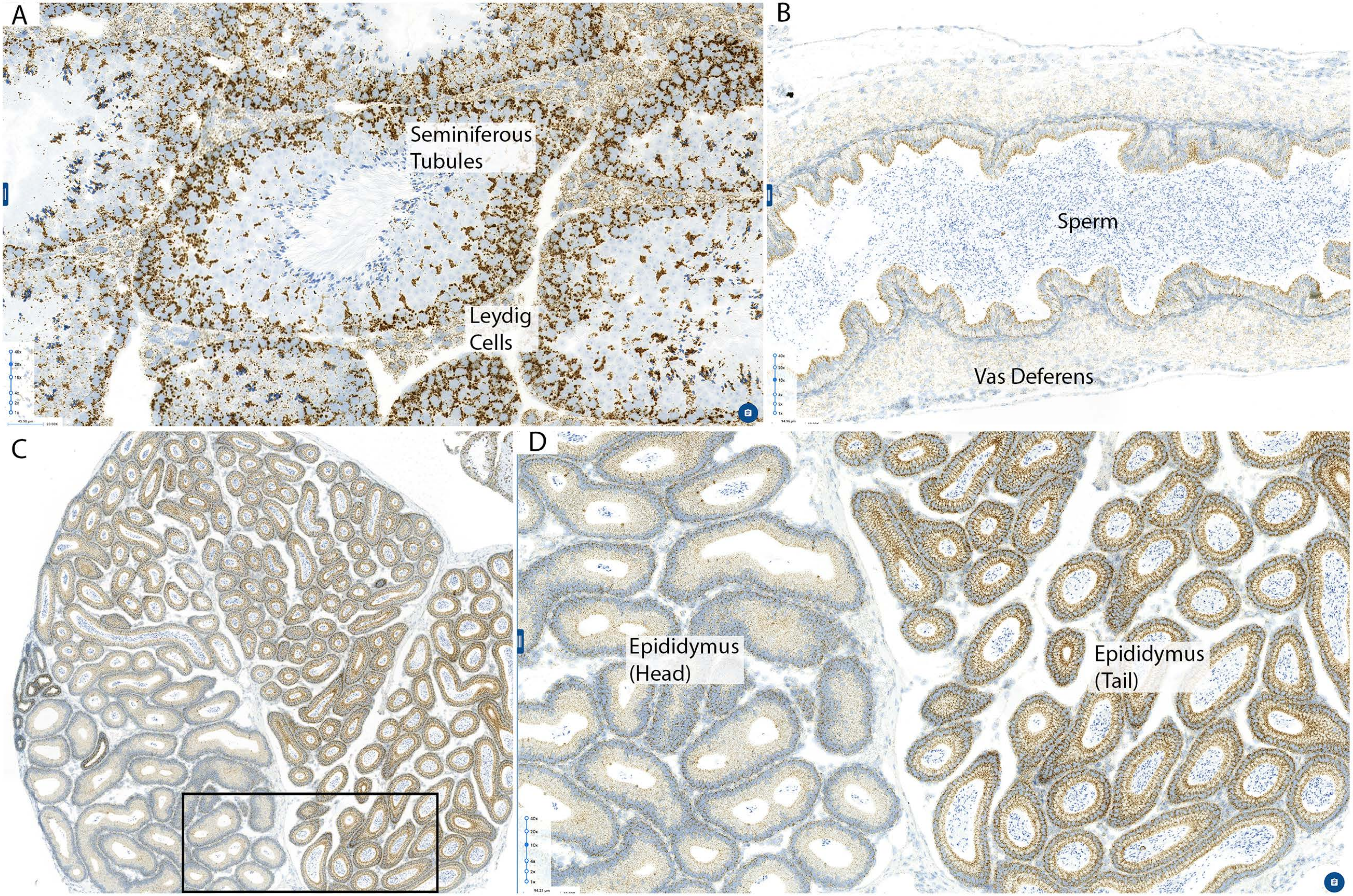
A
Seminiferous
Tubules
Leydig
Cells
B
Sperm
Vas Deferens
C
D
Epididymus
(Head)
Epididymus
(Tail)

### Figure S10

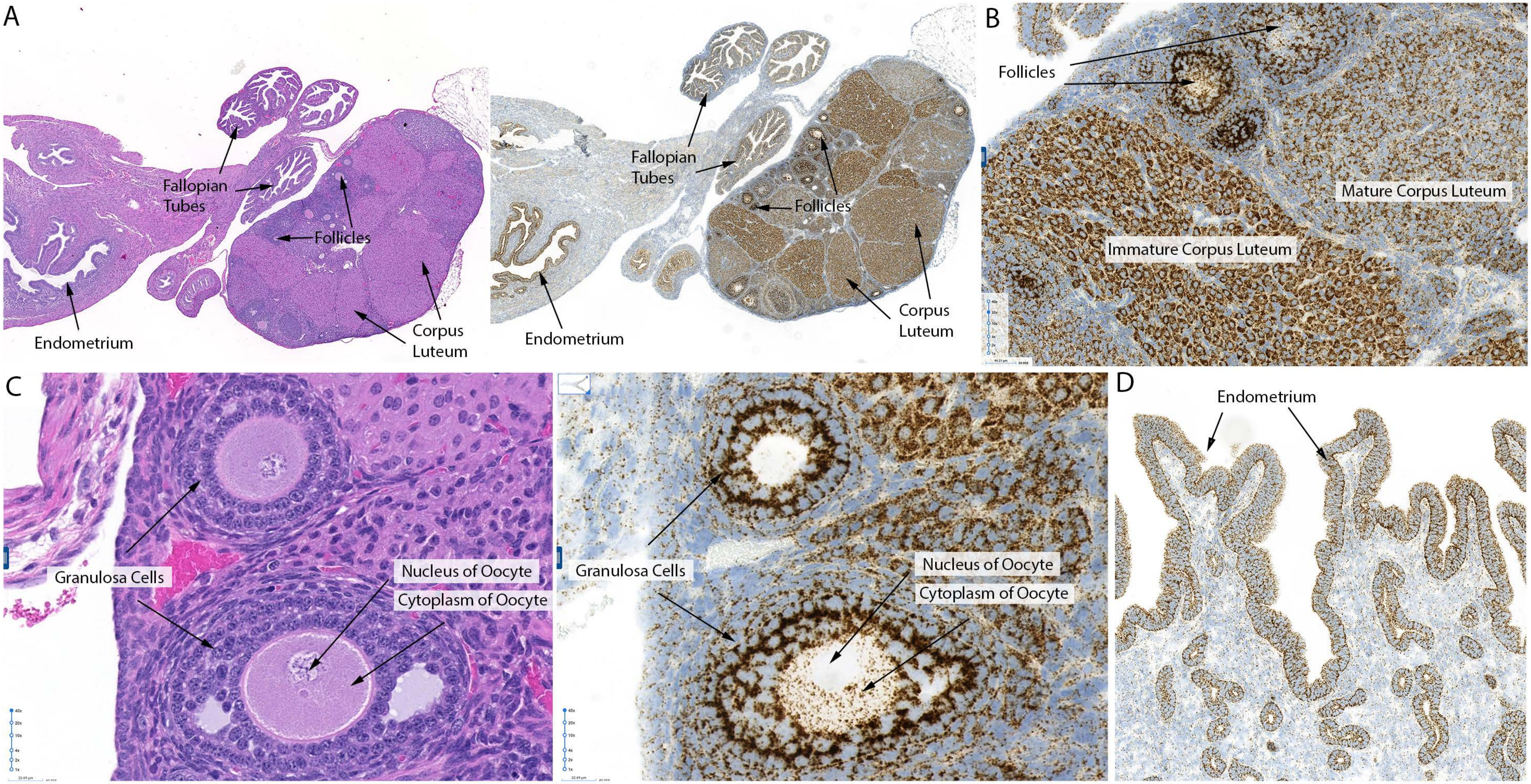
A
Fallopian Tubes
Follicles
Endometrium
Corpus Luteum
Fallopian Tubes
Follicles
Endometrium
Corpus Luteum
B
Follicles
Mature Corpus Luteum
Immature Corpus Luteum
C
Granulosa Cells
Nucleus of Oocyte
Cytoplasm of Oocyte
Granulosa Cells
Nucleus of Oocyte
Cytoplasm of Oocyte
D
Endometrium

### Figure S11

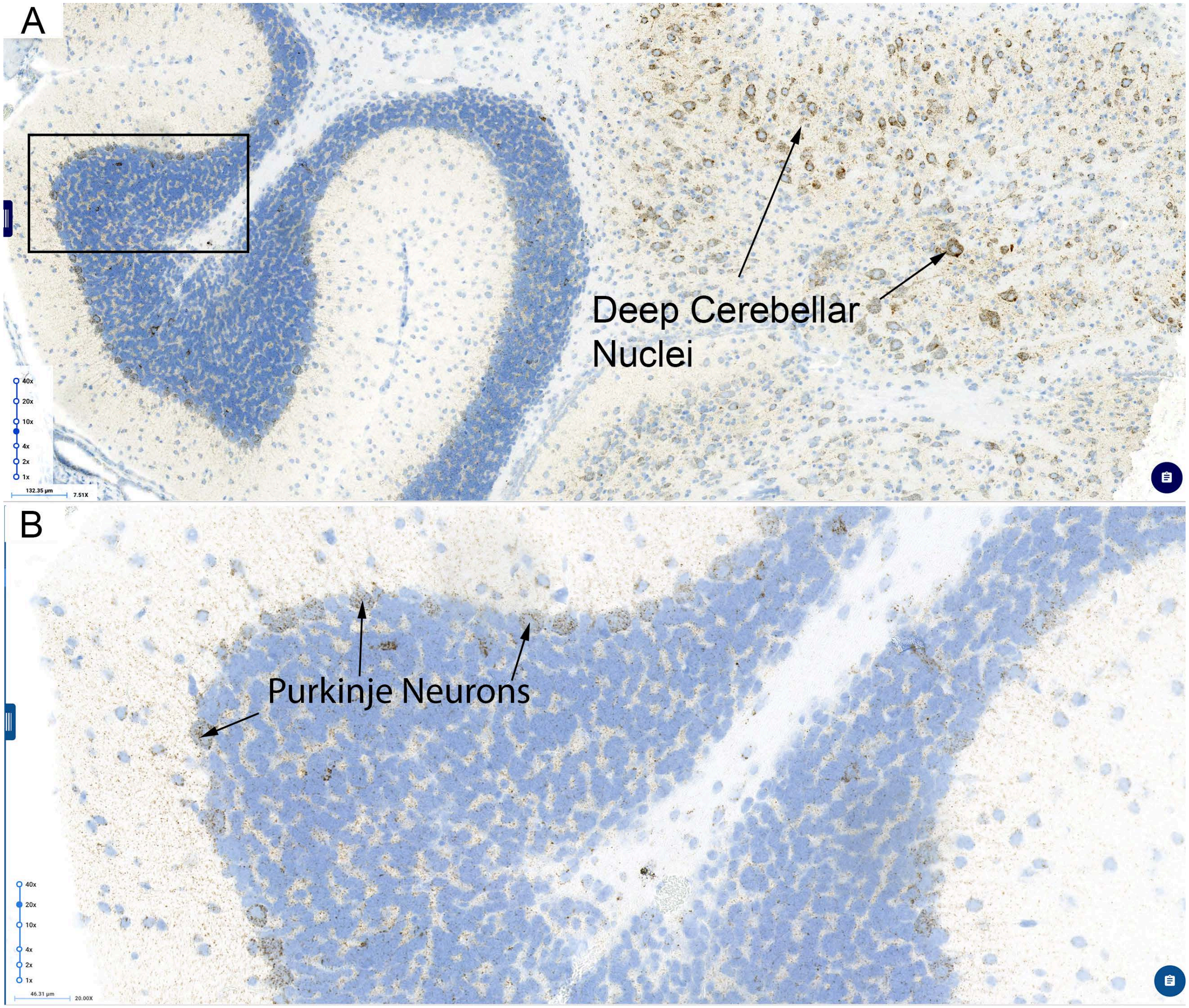
